# The SARS-CoV-2 envelope (E) protein forms a calcium- and voltage-activated calcium channel

**DOI:** 10.1101/2022.10.11.511775

**Authors:** Lysbeth H. Antonides, Quenton W. Hurst, Callum M. Ives, Kiefer Ramberg, Nikitas Ostrovitsa, Eoin Scanlan, Martin Caffrey, Samantha J. Pitt, Ulrich Zachariae

## Abstract

The function of ion channels is essential in the infectious cycle of many viruses. To facilitate viral uptake, maturation and export, viruses must modify the ionic balance of their host cells, in particular of calcium ions (Ca^2+^). Viroporins encoded in the viral genome play a key part in altering the cell’s ionic homeostasis. In SARS-Coronavirus-2 (SARS-CoV-2) – the causative agent of Covid-19 – the envelope (E) protein is considered to form ion channels in ERGIC organellar membranes, whose function is closely linked to disease progression and lethality. Deletion, blockade, or loss-of-function mutation of coronaviral E proteins results in propagation-deficient or attenuated virus variants. The exact physiological function of the E protein, however, is not sufficiently understood. Since one of the key features of the ER is its function as a Ca^2+^ storage compartment, we investigated the activity of E in the context of this cation. Molecular dynamics simulations and voltage-clamp electrophysiological measurements show that E exhibits ion channel activity that is regulated by increased luminal Ca^2+^ concentration, membrane voltage, post-translational protein modification, and negatively charged ERGIC lipids. Particularly, calcium ions bind to a distinct region at the ER-luminal channel entrance, where they activate the channel and maintain the pore in an open state. Also, alongside monovalent ions, the E protein is highly permeable to Ca^2+^. Our results suggest that the physiological role of the E protein is the release of Ca^2+^ from the ER, and that the distinct Ca^2+^ activation site may serve as a promising target for channel blockers, potentially inhibiting the infectious cycle of coronaviruses.

## Introduction

The outbreak of the novel coronavirus disease COVID-19, caused by the coronavirus SARS-CoV-2, has given rise to a pandemic on a global scale. After an extensive vaccination campaign in developed countries over the past 3 years, it is still not under control. New infection waves have continued to arise owing to the emergence of new viral mutants. Currently, new sub-forms of the now dominant SARS-CoV-2 omicron variant continue to present, and with them, the possibility of further variants with increased infectivity and morbidity (1–4). With eradication seemingly beyond our current capabilities, further fundamental knowledge of this pathogen is vital to enable innovation in COVID-19 mitigation.

Coronaviruses (CoVs) mostly cause enzootic infections in mammals and birds. On several occasions, however, they have crossed the species barrier. Notable examples include the severe acute respiratory syndrome virus (SARS-CoV) outbreak in 2003 and the Middle Eastern respiratory syndrome virus (MERS-CoV) outbreak in 2012. Due to the abundant reservoir of zoonotic coronaviruses, it cannot be excluded that further zoonotic coronaviruses will become transmissible to humans in the future. In humans, these viruses can cause upper and lower respiratory tract infections as well as severe acute respiratory syndrome. SARS-CoV-2 is an enveloped positive-strand RNA virus, which encodes 29 proteins including the spike (S), membrane (M) and envelope (E) proteins (5). The S, M and E proteins possess hydrophobic transmembrane domains which embed into the lipid bilayer envelope of SARS-CoV-2. To date, the soluble cytoplasmic domain of S has been the main target in vaccine design (6–8). Meanwhile, the SARS-CoV-2 main protease (M^pro^) has been the focus in the design of small molecule antivirals (9). Inhibitors of the nucleocapsid protein (N) and E have also been proposed (10, 11). Of the four structural proteins, E is the least well-characterized in terms of structure and function. Models of the E protein derived from biophysical data and molecular dynamics (MD) simulations indicate that this 75-residue protein embeds into the lipid viral envelope as a pentamer, comprised of an a-helical transmembrane (TM) domain flanked by less-ordered N- and C-terminal domains on the luminal and cytosolic side, respectively (12–14) (Fig. 1A; the section highlighted in blue and red is the transmembrane region, Fig. 1B). The E pentamer is proposed to function as a viroporin ion channel and is implicated in the assembly and release of viral particles (13–17). The E protein has therefore been put forward as a target for viral neutralization. The rapid mutation rate of the SARS-CoV-2 proteins represents a formidable problem for the efficacy of vaccines and antiviral drugs (2, 18–20). Unlike S, which continues to undergo significant mutations as new variants emerge, E shows a remarkably low propensity to mutate (21) and can therefore be considered a prime target for developing antiviral therapeutics with longer-term efficacy against SARS-CoV-2 and, potentially, future pathogenic coronaviruses (22). The validity of E as a target for biomedical research is highlighted by the prevalence of other envelope proteins with high sequence homology across other related CoVs (14, 21, 22). For example, the E proteins of SARS-CoV and SARS-CoV-2 possess a 95% sequence similarity.

**Fig 1:**
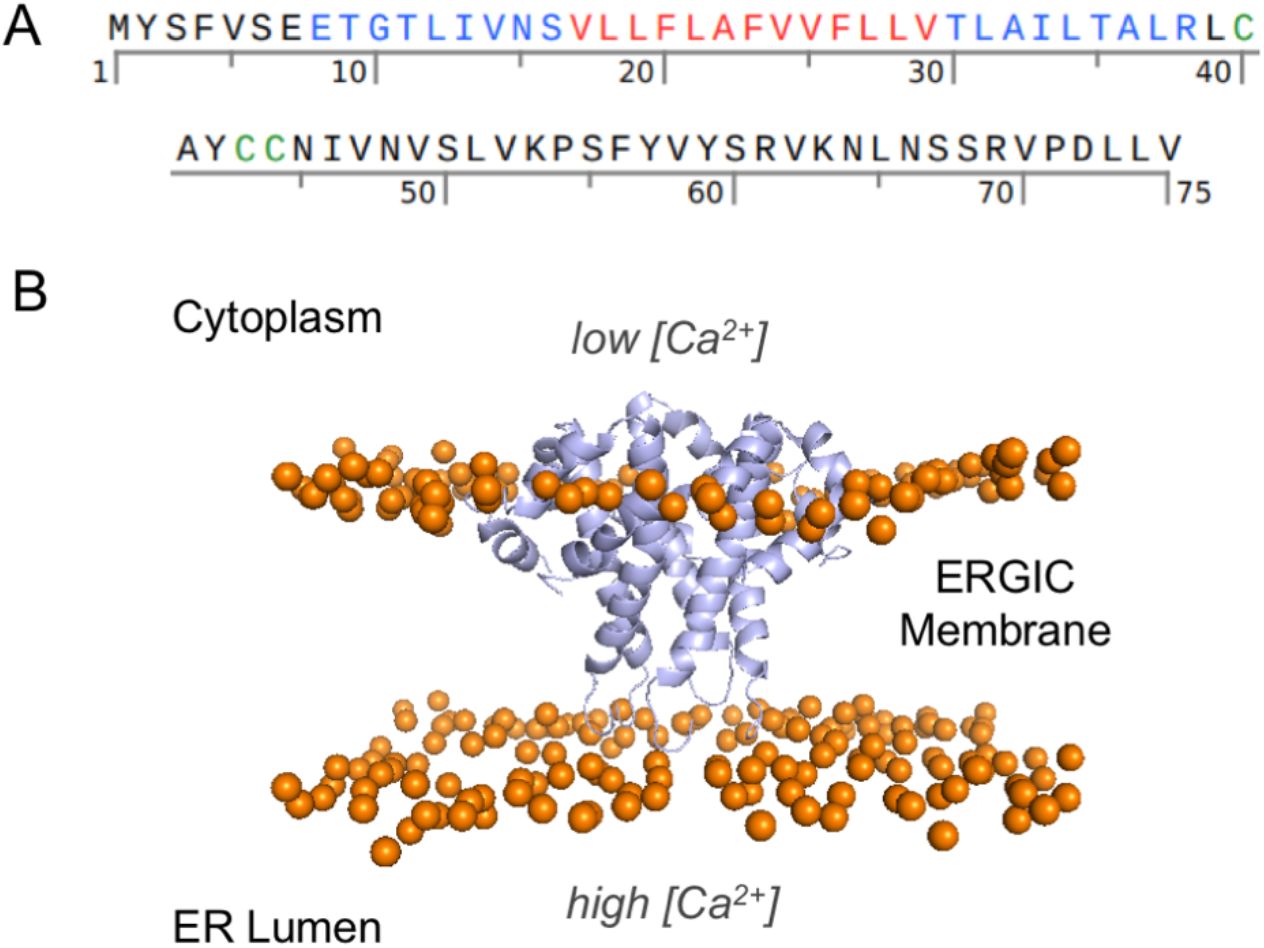
Sequence and structure of the SARS-CoV-2 E protein. (A) Sequence of the E protein. The sections highlighted in blue and red denote the transmembrane region with the section highlighted in red marking the hydrophobic motif. Residues highlighted in green are subject to post-translational modification. **(B)** Side-view of the E protein embedded in the membrane, generated by using Pymol. The lipid headgroup phosphorus atoms are shown in orange. Lipid tails are omitted for clarity.

Viruses commonly alter the ionic homeostasis of host systems to facilitate replication, with calcium ions (Ca^2+^) playing an essential role. In order to alter intracellular ion concentrations, viruses make use of endogenous ion channels of host cells as well as their own channel proteins such as the E protein of SARS-CoV-2. One of the first virus-induced changes in infected host cells is the increase of cytosolic Ca^2+^ concentration, via the influx of extracellular Ca^2+^, as well as a depletion of intracellular Ca^2+^ stores in the ER (Endoplasmic Reticulum), Golgi complex, and lysosomes (17, 23–25). Calcium is proposed to activate transcription factors and enzymes that support viral replication. In addition, Ca^2+^ flow between the ER and the mitochondria is modulated, delaying apoptosis in the first stages of the viral cycle thereby promoting viral replication, and inducing apoptosis in the later stages to assist in the release of new virus particles. During replication of SARS-CoV-2, E is enlisted to modulate intracellular Ca^2+^ stores and likely promotes the formation of virus-like particles (VLPs) by inducing membrane curvature; both processes are necessary for viral budding (12, 17, 23). Inhibition, deletion, or the generation of nonfunctional E protein mutants yield propagation-defective coronavirus variants of the SARS and MERS viruses with drastically reduced viral titer (12, 24–26).

In its physiological setting, the E protein from SARS-CoV channels calcium ions, and other cations and anions (24–27). Importantly, the E protein is responsible for overstimulating inflammatory pathways (25, 27–29) and the so-called ‘cytokine storm’ leading to death in many patients (25, 27). This over-activation of the inflammasome has been linked to the disruption of normal calcium levels in the cell caused by the E protein (25, 27). The permeability of SARS-CoV E to Ca^2+^, as well as its ion selectivity and conductance, are strongly dependent on the composition of the surrounding membrane and the pH of the bathing aqueous solution (29). In recent studies, the E protein from SARS-CoV-2 has also been identified as an ion channel, both computationally (13–17) as well as experimentally (17, 30–32).

Due to the role of the analogous SARS-CoV E protein (25) and the fact that Ca^2+^ is an abundant ion within the ERGIC (Fig. 1B), we hypothesized that the main physiological function of SARS-CoV-2 E upon viral infection is to act as a Ca^2+^ channel, even though the channel is more permeable to monovalent cations under certain membrane conditions (14, 30). We therefore investigated its functional, membrane-bound structure and ion channel activity, focusing on its interaction with Ca^2+^, the surrounding membrane, and the effect of key post-translational modifications. We reveal that the E protein forms a hydrophobically gated ion channel that is activated by Ca^2+^, pH, and electrochemical gradients and that it efficiently conducts Ca^2+^ as well as monovalent ions. The E channel displays a distinct calcium binding region which serves as a recruitment site for ions and an activation region in the pore. In our MD simulations, the channel current shows a strong dependence on the identity of the surrounding lipids and sites that are post-translationally modified with fatty acids. Our findings highlight new ion and lipid interaction regions on the E protein that can serve as targeting sites for designing therapeutic inhibitors of the E protein, potentially averting fatal overstimulation of the host’s immune response and addressing the least mutation-prone component of the SARS-CoV-2 viral proteome.

## Results and Discussion

### SARS-CoV-2 E forms a Ca^2+^-permeable channel in planar lipid bilayers

To investigate the biophysical characteristics of the SARS-CoV-2 E protein we produced and purified a recombinant E construct consisting of the full-length E sequence (residues 1-75) with a C-terminal His10 tag (Fig. S1) (EFL). The same construct was used previously by Hutchinson *et al*. (32) in their study of E localization in eukaryotic cells and displayed ion channel activity. EFL was purified from *E. coli* inclusion bodies into CHAPS micelles for subsequent reconstitution into phosphatidylethanolamine (PE) artificial membranes under voltage-clamp conditions. EFL purification in fos-choline-16 (FC16) detergent was also achieved to high purity (Fig. S2). EFL purified in CHAPS ran predominantly as a monomer (~11 kDa) on SDS PAGE gels (Figure 2A, inset left). Faint higher molecular weight bands are present on the gel consistent with the formation of EFL oligomers and confirmed by Western blot with an anti-His tag antibody (Fig. 2A, inset right). A symmetric main peak was observed in analytical gel filtration profiles of EFL stocks indicating sample homogeneity (Fig. 2A).

**Fig 2:**
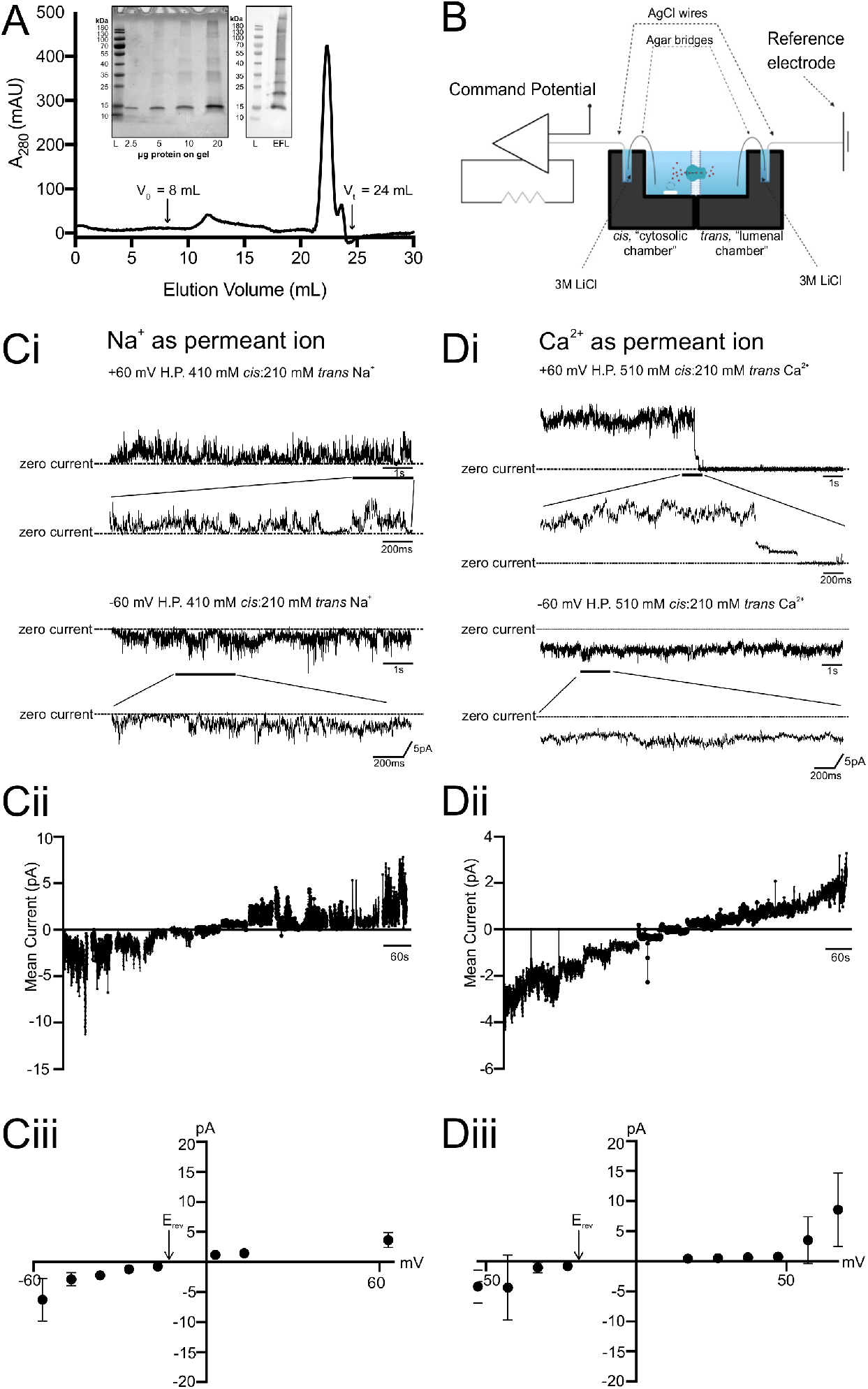
The SARS-CoV-2 E viroporin conducts monovalent and divalent cations. **(A)** Purification of the EFL 10x His construct used for reconstitution into planar lipid bilayers. Analytical gel filtration profile for EFL purified in CHAPS buffer (20 mM TRIS-HCl pH 7.8, 140 mM NaCl, 1 % (w/v) CHAPS, 1 mM DTT). Gel filtration column (Superose® 6 10/300 GL, GE Healthcare) void (V0) and total (Vt) volumes are indicated. Inset left: SDS PAGE (Coomassie stain) loading series. Inset right: SDS PAGE (Western blot with anti-His antibody). **(B)** Schematic of the planar lipid bilayer system used for electrophysiological recordings. Agar bridges are filled with 2 % agarose and 3 M LiCl. All other details are shown in the figure. For more information see Methods. Created with biorender.com. **(C, D)** Typical current recordings of the SARS-CoV2-E (EFL) protein embedded in PE with Na^+^ **(C)** or Ca^2+^ **(D)** as the permeant ion with resultant current-voltage relationships. **(Ci, Di)** The SARS-CoV-2 E protein was incorporated into PE lipids and current fluctuations recorded under voltage-clamp conditions. Recordings are shown at holding potentials +60 mV and – 60 mV and displayed on a 1 s and 200 ms timescale. **(Cii, Dii)** Representative mean current noise analysis plots. Mean current (pA) over time (s) is plotted as a function of voltage (mV) from –60 mV to + 60 mV (with imposed junction potential correction –56.9 mV to +63.1 mV with Na^+^ as the permeant ion and –52.8 mV to 67.2 mV with Ca^2+^ as the permeant ion) in 10 mV steps from left to right. E_rev_ is indicated on the plot. **(Ciii, Diii)** Current-voltage relationship of the SARS CoV-2 E protein constructed from the average mean current of ≥ 3 experiments. Where n ≥ 3, error bars (sd) are shown. Na^+^ conductance was 65.99 ± 15.41 pS (n = 4) and Na^+^ E_rev_ (410 Na^+^ *cis* : 210 Na^+^ *trans*) was –11.6 ± 8.54 mV (n = 4). Ca^2+^ conductance was 11.77 ± 6.87 pS (n = 3) and Ca^2+^ E_rev_ was −18.7 ± 14.52 mV (n = 3; 510 Ca^2+^: 210 Ca^2+^ *trans*). All values shown are corrected for junction potentials.

The oligomeric state of EFL was investigated by mass photometry (MP), a burgeoning technique for mass determination of proteins which has recently begun to be applied to membrane protein targets (33, 34). MP analysis of samples of EFL purified in CHAPS micelles revealed the presence of two distinct species with molecular masses of 64-67 and 188-192 kDa, respectively (Fig. S3). An EFL pentamer would be ~56.7 kDa in size. Complexation of pentameric EFL with 6.15 kDa CHAPS micelles would yield species of ~63 kDa, in line with the MP-determined molecular mass of 64-67 kDa for the most abundant EFL species. The less-abundant larger species is likely composed of multiple EFL oligomers associated with one or more CHAPS micelles. At 188-192 kDa, this species may correspond to a complex of three EFL pentamers and multiple CHAPS micelles. Similar results were obtained from MP experiments with EFL in FC16. Here the molecular mass for the major species ranged from 124 to 143 kDa, consistent with EFL pentamers in FC16 micelles. In line with the MP measurements for CHAPS-solubilized protein, larger species were also detected broaching the possibility that multiple EFL oligomers may occupy single FC16 micelles. Pentamerization of EFL was further confirmed by size exclusion chromatography coupled to multi angle light scattering (SEC-MALS). EFL reconstituted into FC16 detergent micelles yields a single symmetric gel filtration peak and mass analysis via the established triple detector method (35) indicated average molar masses of 145.0 ± 19.5 kDa and 60.4 ± 8.1 kDa for the protein-detergent complex and protein alone, respectively (Fig. S4). These values are in the expected range for an EFL pentamer alone (~56.7 kDa) and in complex with an FC16 micelle (~130 kDa). Protein-micelle complexes up to ~187 kDa in size were detected which may be the result of two EFL pentamers (~113 kDa) occupying one FC16 micelle (~73 kDa). Taken together, the MP and SEC-MALS data indicate that an EFL pentamer is the prevailing state of this construct with lower levels of larger species also occurring.

Channel activity was quantified using voltage-clamp electrophysiological measurements. The experimental arrangement is illustrated in Fig. 2B. Representative examples of activity profiles for bilayer-reconstituted EFL in asymmetric solutions of 410 mM NaCl (*cis* chamber) and 210 mM NaCl (*trans* chamber) at pH 7.2 are shown in Fig. 2Ci. Most reconstitutions gave rise to the apparent insertion of multiple channels with gating to multiple open state levels. Current fluctuations were never observed using elution buffer only (Fig. S5). Using noise analysis, we calculated the mean current at a given voltage and constructed a current-voltage relationship (Fig. 2Cii). Under these conditions, EFL has a conductance of 65.99 ± 15.41 pS (*n* = 4). This is in line with previous measurements made with SARS-CoV-2 E (30). Conductance varies considerably with lipid type and ionic concentrations (30). As observed by Wilson *et al*. (26) with the SARS-CoV E protein, SARS-CoV-2 E displays rectification at the extremities of the voltage ramp. This matches the data of recent work with SARS-CoV-2 E where the protein exhibits outward rectification (31, 36) (Fig. 2Ci-iii). Calculation of the reversal potential (E_rev_) under these conditions yielded values of –11.6 ± 8.54 mV (*n* = 4) (Fig. 2Ciii). This is as expected according to the Nernst equation (−16.89 mV), suggesting that the E protein displays preferred selectivity for cations over anions, with some anion permeation allowed. According to the Goldman-Hodgkin-Katz equation, the recorded reversal potential indicates a permeability for Cl^-^ of about one-third that for Na^+^.

Since the E protein is proposed to form a viroporin that straddles the ERGIC membrane, we sought to further understand its potential physiological role. As Ca^2+^ is an essential cation for ER function, we examined the conductance and relative permeability of Ca^2+^ through the E channel. Figure 2Di shows that the full-length recombinant SARS-CoV-2 E (EFL) protein displays Ca^2+^ permeability (510 mM CaCl_2_ *cis* and 210 mM CaCl_2_ *trans*). Ca^2+^ conductance under these conditions was 11.77 ± 6.87 pS and E_rev_ was −18.7 ± 14.52 *n* = 3 (Fig. 2Dii&iii). Using Ca^2+^ as the permeant ion, we observed incorporation of multiple channels and frequent brief open events to multiple open states. We next assessed the relative Ca^2+^/Na^+^ permeability ratio (P_Ca2+_/P_Na+_) for EFL. The reversal potential was calculated to be –5.54 ± 5.31 mV (*n* = 3) (Fig. 2Diii). Using the Fatt-Ginsborg equation, the P_Ca2+_/P_Na+_ for EFL was calculated at approximately 0.5 ± 0.1 (*n* = 3), suggesting that the viroporin is slightly more permeable to Na^+^ than it is to Ca^2+^ under current experimental conditions.

### Hydrophobic gating and lipid-dependence of SARS-CoV-2 E channels

To gain further insight into the function of the E channel at the atomistic level using MD simulations, we generated a structural model of the Epentamer (residues 8-65), based on the most open conformation of the solution NMR structure ensemble of SARS-CoV E (PDB 5X29, (37)). *In silico* electrophysiology simulations of the channel in membranes were performed to parallel our voltage-clamp experiments with the EFL construct. The model was mutated at five amino acid positions to account for differences in the sequences of the SARS-CoV E NMR construct and wild-type SARS-CoV-2 E. The pentamer was inserted into fully atomistic model membranes, either ERGIC-mimetic or POPC membranes (for details, see Methods). The surrounding aqueous solutions contained either NaCl or CaCl_2_. Initially, 13 MD simulations of 200 ns length were performed to construct and equilibrate the molecular models of the E protein embedded in the lipid bilayers.

In all simulations, the E channel was fully hydrated prior to equilibration. However, during equilibration, most channels showed hydrophobic dewetting around the main hydrophobic motif (highlighted in red in Fig. 1A), in accordance with earlier observations (Fig. S6; (38)). Reversible dewetting is an established channel regulatory mechanism for viroporins (39), and the dewetted state of the pore is unable to conduct ions. However, ERGIC and other organellar membranes experience strong electrochemical gradients that may facilitate what is referred to as electrowetting. This is analogous to the electrowetting process observed in artificial hydrophobic carbon nanopores where rehydration of dewetted pores happens in response to an applied voltage (40). Testing the behavior of the E channel under voltage, we found that the application of an electric field can indeed rehydrate the pore, which in turn facilitates ion permeation. This is true for simulations performed with both Na^+^ and Ca^2+^ ions. Along with the presence of specific voltage and ion gradients across the ERGIC membrane, our simulations indicate that the E protein is likely to be a voltage-gated pore that is regulated by the hydrophobic gating mechanism and electrowetting (39, 40). Figure 3A-C show the Ca^2+^ ion permeation events, the number of water molecules residing within the pore, and the pore radius in a set of simulations of the E protein in an ERGIC-mimetic membrane surrounded by a Ca^2+^ solution. As can be seen, the pore radius and hydration levels are intrinsically linked. The cyan arrow and rectangles in Fig 3A-C denote times at which a decrease in pore radius and hydration was detected, coupled to a cessation of permeation events (e.g., after 125 ns of simulated time in CaCl_2_ replicate simulation 2). A pore hydration level below ~70 water molecules cannot sustain ion permeation. This phenomenon is even more pronounced in the systems simulated with Na^+^ ions (Fig. S7). Notably, ion conduction continues after the pore diameter, and thereby pore hydration increases again.

**Fig 3:**
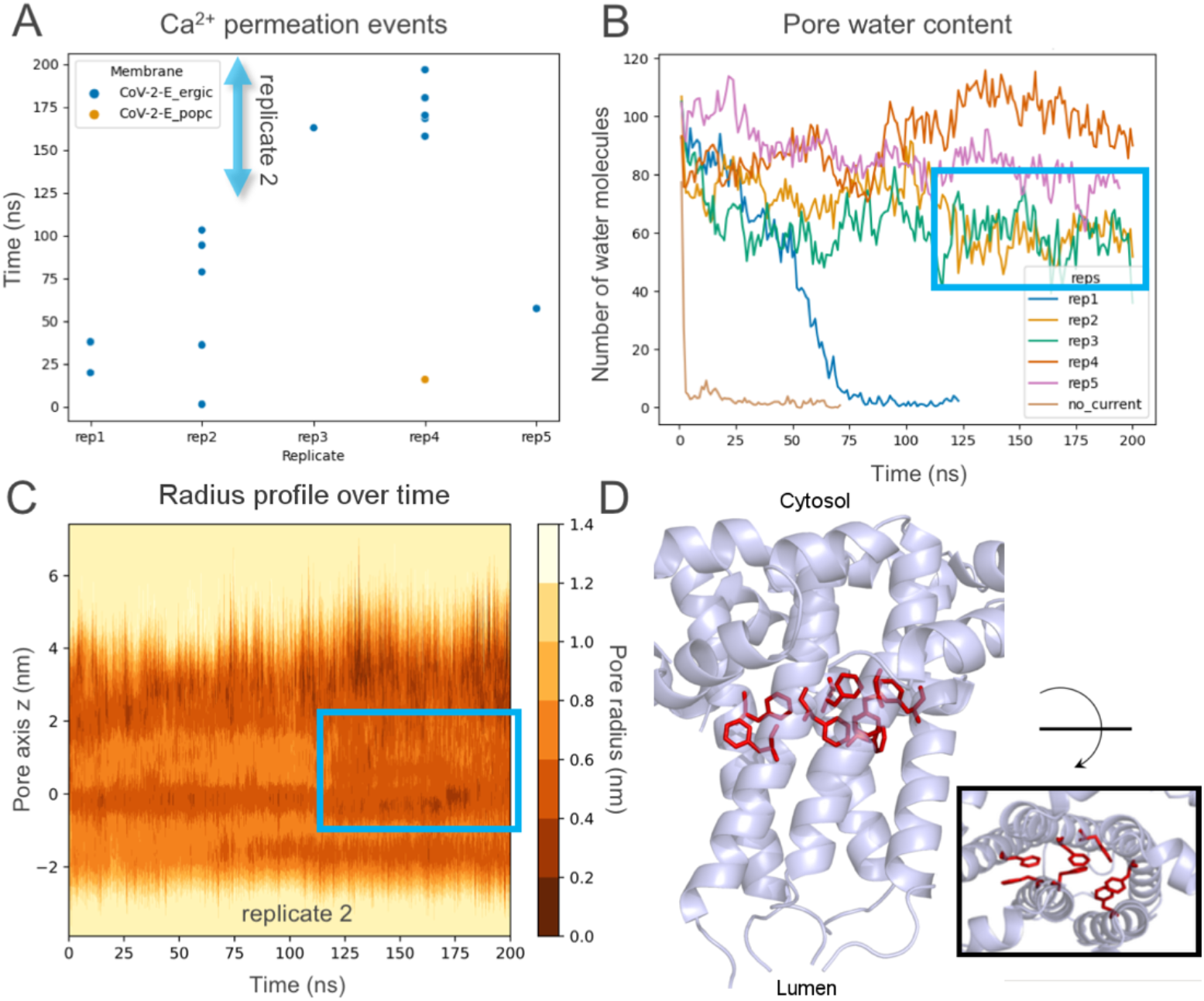
Ca^2+^ permeation events and the link between pore radius and water content. **(A)** Permeation events of Ca^2+^ through the E protein embedded in ERGIC-mimetic membranes and POPC membranes, shown in blue and orange, respectively. Five replicates of each setup were conducted. The cyan arrow indicates the time period in replicate simulation 2 where the pore radius became too narrow to facilitate ion permeation. **(B)** Graph showing the number of water molecules occupying the transmembrane region of the pore. The cyan rectangle highlights the decrease in water molecules in replicate 2, which is caused by a decrease in pore radius. Water was excluded from the pore nearly instantly in the simulation without an applied voltage (no-current). **(C)** Plot showing the radius profile of the pore in replicate simulation 2 throughout the simulation. Darker colors denote a smaller pore radius. The cyan rectangle shows the decrease in pore radius which causes the decrease in pore water content shown in (B) and the halt of permeation shown in (A). **(D)** Structure of the E protein showing the collapse of the Phe residues in the pore hydrophobic gate, closing the channel (side view; inset: top view).

In the center of the pore, a hydrophobic gating motif (consisting of Phe20, Phe23 and Phe26) acts as a further regulatory element. The areas highlighted in red in Fig. S5 illustrate the total collapse of the pore, where the radius around the hydrophobic motif nears zero, which is followed by complete pore dewetting. When the phenylalanine side chains from the 5 protomers point into the center of the pore, following a closing movement caused by Phe-Phe hydrophobic interactions from different subunits (Fig. 3D), the water molecules are expelled from the pore even in the presence of an applied voltage. Similar models of the E TM domain in Phe-regulated ‘open’ and ‘closed’ states have been put forward by the Hong lab, informed by solid state NMR experiments with a TM construct (residues 8-38) reconstituted into lipid bilayers (41). Furthermore, the conformation of the phenylalanine residues is affected by the acyl chains of neighboring phospholipids and by post-translational modifications, as described below.

There is a clear difference in pore stability and openness between simulations in ERGIC and POPC membranes. In ERGIC membranes, all except one replicate simulation display an open, conductive pore, whereas in systems with a simple POPC model membrane, dewetting occurs in all but one replicate. This is also the only replicate that allows an ion to permeate in POPC, as indicated by the orange circle in Fig. 3A. The flow of hydrated ions facilitates the transport of water molecules through the pore, which aids in preserving an open, hydrated state. These observations serve to highlight how the membrane environment affects E protein function. Channel openness and ion conduction appear to be enhanced by the phospholipid environment commonly found in ERGIC organellar membranes, in line with the physiological localization of the CoV-2-E protein (12, 32). These findings are in agreement with previous experimental studies of the E protein, which established a strong lipid dependence of SARS-CoV E currents, with a particularly important role played by negatively charged phospholipid molecules in stabilizing the pore (25). Our simulations suggest that negatively charged lipids in ERGIC-mimetic membranes, especially POPI, form intimate contacts with the Glu8 residues on the luminal side of the membrane-facing surface of the E protein. In doing so, a functionally relevant cation chelating or binding site is formed over most of the duration of our simulations (Fig. 4A; Table S1; for further details see below).

**Fig 4:**
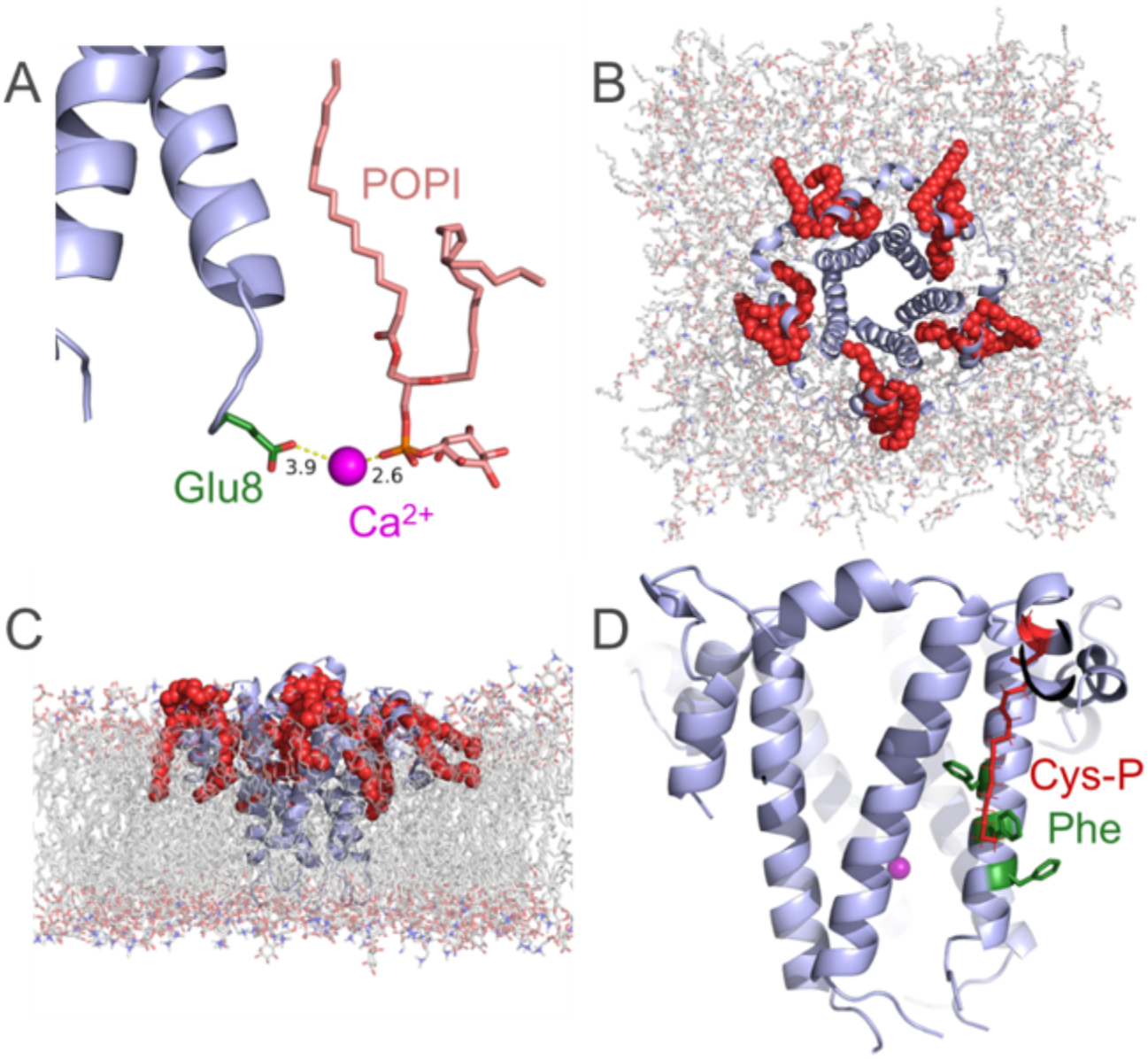
MD simulations of the interactions of SARS-CoV-2 E with lipid headgroups and its palmitoyl post-translational modifications. **(A)** Structure of the luminal *N*-terminus of the E protein, showing the chelation of a Ca^2+^ ion by the Glu8 side chain and the negatively charged headgroup of POPI. **(B)** Top view from the lumen of the E protein embedded in an ERGIC-mimetic membrane. Highlighted in red are the three palmitoylated cysteine residues per monomer, 15 in total. **(C)** Side view of the E protein in the membrane. The palmitoyl chains are embedded in the lipid bilayer and serve to anchor the E protein in the membrane. **(D)** Structure of the E protein in the membrane with membrane phospholipids hidden. Highlighted in red is a palmitoylated cysteine residue, Cys-P, and in green are the three phenylalanine residues that are part of the hydrophobic motif. The interaction between the palmitoyl moiety in Cys-P and two of the phenylalanine residues stabilizes the pore in an open state by favoring the phenyl rings to orient into the membrane and away from the center of the pore.

### The effect of post-translational modification of the SARS-CoV-2 E protein

It is well-known that coronaviral proteins, including those from SARS-CoV-2, often carry a number of post-translational modifications (PTMs; see, e.g., (42)). A common PTM of cysteine residues is S-palmitoylation, due to the functionalization of the sulfur atom in these residues (12, 16, 42–44). The SARS-CoV E protein is known to be palmitoylated, and experimental evidence suggests that all three cysteine residues adjacent to the TM domain serve as targets for palmitoylation (12). For other coronaviruses, triple-cysteine mutated E proteins, which cannot be palmitoylated, have been shown to be unstable, prone to degradation, and to significantly reduce the viral yield. Lopez *et al*. (43) rationalize this loss of stability by suggesting that palmitoylation of the coronaviral E proteins might affect virus-host membrane interactions with the loss of the membrane-anchoring lipid chains leading to decreased membrane stability (16, 43–45). Furthermore, the addition of a hydrophobic tail is known to aid in the trafficking and subcellular localization of the E protein (16).

Considering the high degree of sequence identity between the SARS-CoV E and SARS-CoV-2 E proteins, we palmitoylated all three cysteine residues (Cys40, Cys43, Cys44) in each subunit in our pentameric model of the SARS-CoV-2 E protein channel (Fig. 4B,C; sequence positions highlighted in green in Fig. 1A). Notably, none of the simulations of systems without palmitoylation produced a stable pore. The addition of palmitoylation of all three cysteines led to a substantial stabilization of the pore, in both POPC and ERGIC-mimetic membranes. Further, we found that the pore open state was stabilized mainly as a result of an interaction between the palmitoyl chain on Cys44 and Phe23 in the hydrophobic gate region of the E protein channel in each of its monomers. During our simulations, at least one palmitoyl chain interacted with the phenylalanine residues over 52.7% of the time in systems with CaCl_2_ and ERGIC membranes, 43.7% of the time in systems with NaCl and ERGIC membranes, and 33.3% of the time in systems with CaCl_2_ and POPC membranes (Tables S2, S3 and S5, respectively). This interaction prevented the aromatic ring of the phenylalanine from associating with the corresponding phenylalanine residues from other monomers inside the channel, which would have resulted in occluding the pore (Fig. 3D). The palmitoyl moiety attached to Cys44, when positioned between two monomers, appears to be the most important of the three modified cysteine residues with regard to stabilizing the channel. In the simulations that display the largest level of channel opening, the palmitoyl chain of Cys44 inserts snugly between individual protein monomers presumably contributing to preventing their collapse (Fig. 4D). By contrast, the attraction of the phenylalanine residues for each other in the center of the pore causes the pore to collapse in the majority of the simulations and especially in the case of unmodified, palmitoyl-free channels. By comparison with Cys44, the palmitoyl chains attached to Cys40 and Cys43 show no obvious interaction with residues of the hydrophobic phenylalanine gating motif, or indeed with the pore. However, our simulations indicate that they act as an anchor which helps to keep the protein in place within the membrane and in functionally-relevant proximity to the other protein monomers (Fig. 4B,C). Our computational results thus show that palmitoylation of at least one cysteine residue promotes a stable and open E protein pore. This was also noted by Sun *et al*. (16), who reported a decrease in pore radius and subsequent collapse of the E pentamer pore when simulating the non-palmitoylated E protein. In the palmitoylated structure, a larger pore radius and enhanced stability was observed. Parenthetically, we note that some of the palmitoyl chains in the Sun *et al*. simulation extended beyond the membrane into the bathing aqueous solution. We consider this unlikely to represent a physiologically relevant conformation for the hydrophobic acyl chains.

### SARS-CoV-2 E exhibits a binding ring for calcium ions at the luminal pore entrance

In our simulations, the open SARS-CoV-2 E protein channel conducts Na^+^ and Ca^2+^ ions toward the negatively polarized side of the bilayer and Cl^-^ ions in the opposite direction. A slightly raised permeability for Cl^-^ was seen in our simulations as compared to our experiments. This finding is consistent with previous experimental electrophysiology recordings on the homologous E protein in which a delicate dependence of the channel’s ion selectivity on its precise environment was observed, tipping the balance toward either a slight preference for anions or cations (23, 25). Notably, we observe that cations on the luminal side interact intimately with the five Glu8 residues that line the luminal entrance to the pore (Fig. 4A, Fig. 5). SARS-CoV-2 E localizes to the ERGIC, connecting the ER and Golgi compartments (31). The pH of these cellular compartments ranges from neutral (7.2) to slightly acidic (6.0) (46). In this pH range, glutamic acid residues are expected to be deprotonated and to carry a formal net charge of −1e. Therefore, a ring with a strongly negative electrostatic potential is formed by the Glu8 residues of the five monomers at the luminal entrance to the TM domain. We observe an aggregation of cations in the vicinity of this glutamate ring in each of the simulations, enhanced further by the negatively charged lipids, POPI in particular, that reside in this region of the protein (Fig. 4A). The most common distance for calcium-oxygen interactions is reported to be in the range 2.5 Å to 3.5 Å (47). Accordingly, we investigated the association of cations within a sphere of radius 3.5 Å around each of the glutamate side chains. We found that both Na^+^ and Ca^2+^ bind to the ring of glutamate residues with a strong preference for Ca^2+^ ions. In the simulations with Ca^2+^, the ring is occupied with one or more Ca^2+^ ions for 61% of the time in ERGIC membranes and 53% of the time in POPC membranes (Tables S5, S6). Our simulations suggest that the difference is due to the additional interactions with POPI, whose phosphate group chelates the divalent Ca^2+^ cations jointly with the negatively charged glutamate side chains (Fig. 4A), stabilizing their binding further. Importantly, this finding suggests a synergistic effect of Ca^2+^ and the presence of negative lipids in maintaining the pore in an open state. With Na^+^, the occupancy of the ring with at least one Na^+^ ion was considerably lower, amounting only to 15% of the time in ERGIC lipid membranes. The residence times for ions at the binding ring were found to be between 1 and 2 ns for Ca^2+^ and about 0.3 ns for Na^+^ (Table S1 and S2 in Supporting Information). A density of Ca^2+^ ions over a 200 ns simulation is shown in Fig. 5, demonstrating the enrichment of Ca^2+^ ions around the luminal glutamate ring, which facilitates ion entry into the pore.

**Fig 5:**
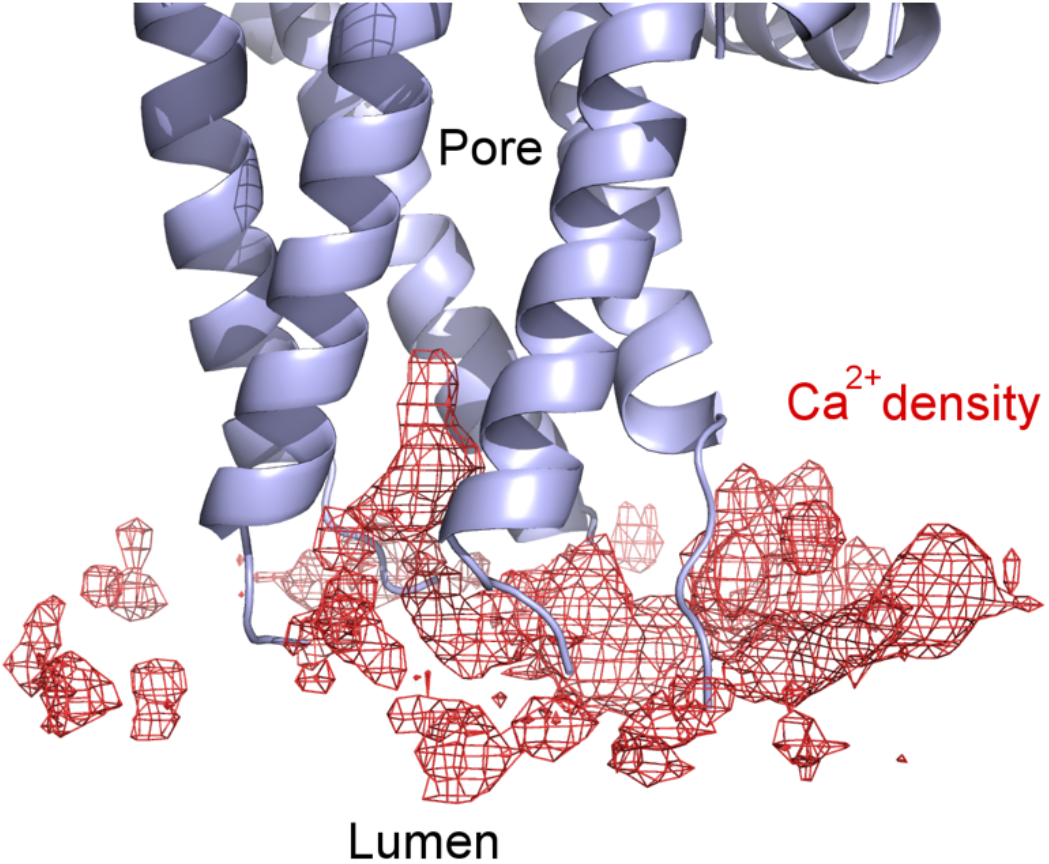
Calcium binding region around the luminal pore entrance of SARS-CoV-2 E. The structure of the E protein is shown with phospholipids removed. The red mesh shows the density of Ca^2+^ ions at the luminal surface and extending into the pore entrance over a 200 ns simulation.

We also tested experimentally the effect of luminal Ca^2+^ on the gating properties of EFL (Fig. 6). Under voltage-clamp conditions, using Na^+^ as the permeant ion, we found that increasing the luminal [Ca^2+^] to 50 μM had no significant effect on the gating properties of EFL (Dunn’s multiple comparisons post-test, control vs. 50 μM CaCl_2_, p=0.2907). In contrast, when the luminal [Ca^2+^] was raised to 100 μM, a significant increase in the average mean current was consistently observed, indicating stabilization of the open state of the channel. Further elevations in the luminal Ca^2+^ concentration to 300-1000 μM increased channel activity in a dose-dependent manner. (Dunn’s multiple comparisons post-tests were run with the following p-value results, control vs. 100 μM CaCl_2_ = 0.0333, control vs. 300 μM CaCl_2_ = 0.0153, control vs. 700 μM CaCl_2_ = 0.0031, control vs. 1 mM CaCl_2_ = 0.0974.) While the variability in channel activity at 1000 μM was too great to determine significance, the trend of increased current at this concentration was also observed. Due to the fast and flickering nature of channel gating and the observation that multiple channels gate in the bilayer, we are unable to make any comment on the dwell times of channel openings.

**Fig 6:**
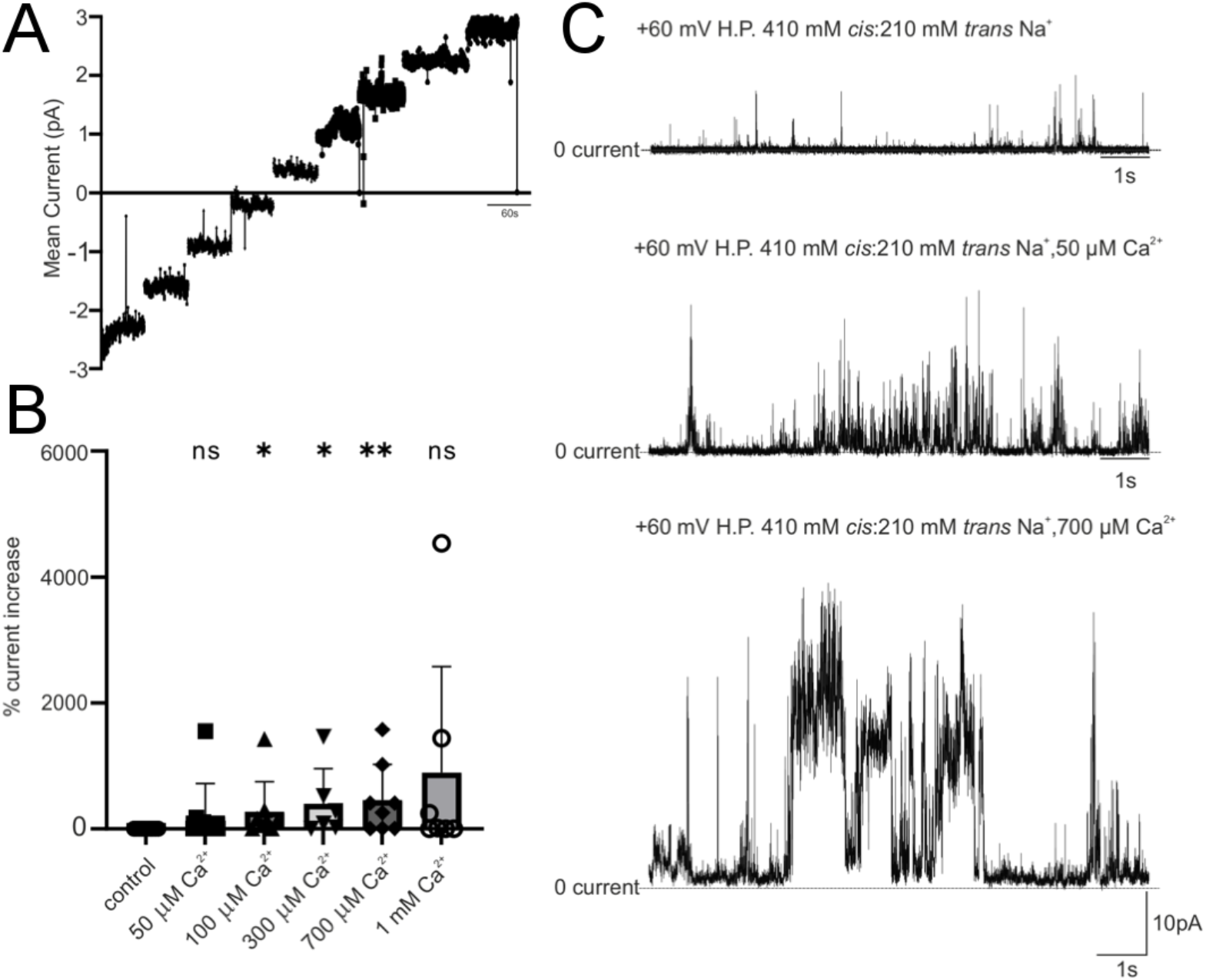
Ca^2+^ regulation of EFL in a PE lipid membrane. **(A)** Representative mean current noise analysis plot under bi-ionic conditions (210 mM Na^+^ *cis*: 210 mM Ca^2+^ *trans*). To determine E_rev_ under these conditions, the mean current in pA over time is plotted as a function of voltage from −4 to 3 mV in 1 mV steps with junction potential correction imposed (−6.2 mV to 1.4 mV). E_rev_ was –5.54 ± 5.31 mV (sd) (n = 3). **(B)** Representative current recordings of EFL with Na^+^ as the permeant ion following subsequent addition of incremental concentrations of luminal (*trans*) Ca^2+^. Recording conditions are indicated above each trace. **(C)** Bar chart showing the combined data from n ≥ 4 recordings. A dose-dependent increase of channel activity with increased luminal Ca^2+^ is observed. Significance was tested by a Kruskal-Wallis one-way non-parametric ANOVA (P ~ 0.0076) followed by Dunn’s post-multiple comparisons test; p-values are as follows: control vs. 50 μM CaCl_2_ = 0.2907, control vs. 100 μM CaCl_2_ = 0.0333, control vs. 300 μM CaCl_2_ = 0.0153, control vs. 700 μM CaCl_2_ = 0.0031, control vs. 1 mM CaCl_2_ = 0.0974.

These findings confirm the results from our simulations and demonstrate that luminal Ca^2+^ ions activate the E-protein channel. A potential explanation could be that increasing local Ca^2+^ concentration facilitates movement of the ions into the channel by mass action. Additionally, the Ca^2+^ interacts with the Glu residues and in doing so, alters the conformation of E, thereby likely favoring channel opening and ion flow into and through the channel.

### Functional studies of the isolated SARS-CoV-2 E transmembrane domain

To date, only the TM domain of SARS-CoV-2 E (ETM) has been structurally characterized (48). We therefore set out to determine if the TM region is a functional substructure of the full-length E protein. Accordingly, we produced ETM, consisting of residues 8-38 of SARS-CoV-2 E (Fig. S1), by solid-phase peptide synthesis (Fig. S8). This construct is analogous to the one characterized by solid state NMR spectroscopy by the Hong lab (48). As with the EFL construct, we tested the functionality of ETM in *in vitro* voltage-clamp experiments. ETM was reconstituted into the same CHAPS-containing buffer used with the EFL construct prior to measurements. We first examined whether ETM was inserted into PE planar lipid bilayers under voltage-clamp conditions. Applying the same experimental conditions used to characterize EFL (410 mM NaCl in the *cis* chamber and 210 mM NaCl in the *trans* chamber, Fig. 1B), we did not observe any channel-like activity for ETM in any of our experiments (n = 7) when the membrane was clamped in the range from −60 mV to +60 mV (Fig. S9). Whereas Wilson *et al*. (26) have demonstrated activity for the SARS-CoV E TM domain along with Torres *et al*. (49) and Verdiá-Báguenaa *et al*. (23), under our experimental conditions and voltage range we do not observe activity for SARS-CoV-2 ETM domain alone. We note that pore activity has been reported for a SARS-CoV-2 ETM construct in *Xenopus* oocytes using electrophysiology experiments (50). This difference in activity may be due to the presence of a wide range of other proteins and channels in the oocyte membranes, which potentially interact with the E protein. Further experiments, involving a broader spectrum of techniques, are required to resolve the discrepancy.

To extend our experimental observations, we next carried out MD simulations on isolated ETM domains in assembled pentameric form. In our previous simulations, the full-length protein was found to be most stable and functional in an ERGIC-mimetic membrane and in solutions containing Ca^2+^ ions. We therefore used the same conditions for our simulations of the TM-constructs. The TM domains of the two published E-protein NMR structures (PDB ID: 7K3G; (48) and PDB ID: 5X29 (residues 8-38), highlighted in blue in Figure 1a; (37)) were simulated. Both were incorporated in ERGIC-mimetic membranes and subjected to MD simulations under voltage-clamp conditions, as above. In all replicates, the closed dewetted conformation alone was observed. Pore formation did not occur, and no ion permeation was observed for either construct (Fig. S10). We detected a small degree of pore hydration in the ERGIC-mimetic membrane system. However, compared to the hydration levels associated with a functional pore, it was much more limited and did not sustain ion permeation under voltage. In a POPC membrane, the pore closed and dewetted instantly, demonstrating once again the sensitivity of the protein’s TM-region to the lipid composition of the surrounding membrane. These findings agree with our results from the *in vitro* experiments with ETM, which indicated that the isolated TM domain does not sustain ion channel current in planar lipid membranes. Taken together, our experimental and computational results suggest that the TM section alone does not form a physiologically functional substructure of the SARS-CoV-2 E protein. TM domain (residues 8-38) constructs may therefore not be suitable models with which to gain physiologically-relevant insights into the structure and function of SARS-CoV-2 E and for drug design and discovery.

## Conclusion

In this study we have shown that the E protein of SARS-CoV-2 forms a viroporin that is regulated by the luminal Ca^2+^ concentration, the presence of electrochemical gradients, such as those existing across the membrane of the ERGIC subcellular compartment (the likely physiological location of the E protein in the host cell), and the negatively charged phospholipids typical of ERGIC membranes. Notably, we demonstrate that the E protein displays a luminal annulus to which Ca^2+^ binds and in so doing stabilizes the open state of the pore. As a consequence, Ca^2+^ ions increase pore open times and, in turn, ionic current through the E channel. We show experimentally that increasing luminal Ca^2+^ concentration to 100 μM consistently increased channel activity, again suggesting that calcium is a key factor for stabilizing the open state of the channel. Importantly, the range of luminal Ca^2+^ concentrations that affects E protein gating (0.1–1 mM) falls within the expected range of luminal Ca^2+^ concentration in the ERGIC (51). We also reveal that SARS CoV-2 E is more selective for cations than anions but that it does not show a strong discrimination between monovalent and divalent cations. The slight preference for monovalent cations seen matches previous reports on E proteins (23, 25). Given its localization in the ERGIC membrane and the large Ca^2+^ gradient that exists across the membrane of this Ca^2+^ storage compartment, we suggest a role for the SARS-CoV-2 E protein in mediating intracellular Ca^2+^ signaling events that are consistent with previous reports on E from SARS-CoV (51). Our data suggests that Ca^2+^ release through the SARS-CoV-2 E viral pore is strongly dependent on the Ca^2+^ load. Depletion of Ca^2+^ stores below a threshold level or modulation of the cation binding site could terminate E protein mediated calcium flux. This is important given the emerging role of intracellular Ca^2+^ dysregulation during SARS-CoV-2 infection (52–55). Our findings therefore indicate that the identified cation binding site could be a suitable target for the design of channel-inhibiting antivirals (56). Importantly, the E protein displays the least sequence variability of all proteins encoded by the SARS-CoV-2 genome, and may therefore represent an especially useful target in the face of the high mutation rate of the virus, which has rapidly given rise to increasingly infectious and vaccination-escaping variants in recent months (14, 21). In addition, the similarity of SARS-CoV-2 E to other zoonotic coronaviruses is high. In the future, these may cross the species barrier to humans. Therefore, drugs targeting the SARS-CoV-2 E channel may serve as a lead for broad-spectrum coronavirus anti-virals.

## Materials and Methods

### Simulation system setup

To systematically study the behavior and function of the SARS-CoV-2 E protein, the 3D structural model of SARS-CoV-2 E was embedded into a lipid bilayer and solvated using CHARMM-GUI (57). The protein was adapted from PDB 5×29 (37) (SARS-CoV-2 E protein) by mutation of the following residues according to the SARS-CoV-2 E sequence: Ala40Cys, Ala43Cys, Ala44Cys, Thr55Ser and Val56Phe. The cysteine residues were palmitoylated using CHARMM-GUI; the palmitoyl chains were then manually adjusted to be accommodated within the lipid bilayer with no clashing atomic contacts. Two systems were generated, one consisting of a membrane of pure POPC lipids. The second was composed of an ERGIC-mimetic membrane, containing 50% POPC, 25% POPE, 10% POPI, 5% POPS and 10% cholesterol. For solvation, the TIP3 water model was used and 150 mM NaCl or CaCl_2_ were added using GROMACS 2021.1 (58). In simulations with calcium, an alternative multi-site model for calcium was used, which was developed by Zhang *et al*. (59). The CHARMM36-mar2019 forcefield was used and the system was subsequently minimized and equilibrated, prior to MD simulation, using the equilibration procedure supplied by the CHARMM-GUI (www.charmm-gui.org) (57). In short, the systems were equilibrated in 1.85 ns, and the force constraints were gradually released over six equilibration steps. During the following 200 ns equilibration simulations, the Nose-Hoover thermostat was used to maintain the temperature at 310 K (60). The Parrinello-Rahman barostat was used to maintain pressure semi-isotropically at 1 bar (61). Periodic boundary conditions were used throughout each simulation. The particle-mesh Ewald method was used to model long-range electrostatic interactions, with a cut-off of 12 Å (62). In order to constrain bonds with hydrogen atoms, the LINCS algorithm was used (63). In total, four systems were generated; system 1 and 2 with POPC and NaCl and CaCl_2_ ions, respectively, and system 3 and 4 with an ERGIC-mimetic membrane and NaCl and CaCl_2_ ions, respectively. Each production simulation was 200 ns long, used the same approaches, and was repeated five times for each system. All simulations were carried out with the simulation package GROMACS 2021.1 (www.gromacs.org) (58).

### Molecular dynamics simulations and *in silico* electrophysiology

In order to develop a stable model of the E protein embedded in a membrane, 13 molecular dynamics simulations of 200 ns were performed. The majority of these were of the protein in an ERGIC-mimetic membrane with either NaCl or CaCl_2_ ions. All systems showed a fully hydrated SARS-CoV-2 E pore prior to equilibration. However, most dewetted around the hydrophobic motif (Residues Phe23 and Phe26) during equilibration. Subsequent MD simulations without a voltage caused the pores to collapse. Any systems which were simulated without a voltage were unstable. The pore collapsed consistently within the first few nanoseconds of the MD simulation. Simulations with an applied voltage or a transmembrane electrochemical gradient rewetted, and many exhibited open pores through to the end of the simulations or closed-open transitions. In our simulations with voltage, we applied a constant membrane potential of 800 mV (64, 65).

The MD trajectories were analyzed using in-house Python scripts, Gromacs utilities and MDAnalysis (66). CHAP (67) was used to analyze the pore structure and radii. Plots were generated with Python, using Matplotlib (68). Ion permeation events were identified using an in-house written python script, and verified visually, using VMD (69). The interactions between lipids and palmitoylated Cys-residues were analyzed using in-house written Python scripts. The analyses were performed on five replicates of 200 ns simulations of each system with a 10 ps time-step. The cation binding site near the glutamate (Glu8) residues were identified as follows. A time series of each permeating ion analogous to its position along the pore axis was plotted. Furthermore, the PENSA *Featurizer* function (70) was used to determine the 3D density maxima of the cations within 3.5 Å of the glutamate residues. In order to calculate the ion occupancy probabilities of the cations at the predetermined glutamate binding sites, N_occupied_, the number of frames wherein an ion’s center of geometry is within 3.5 Å of the center of geometry of the ion coordinating binding site (in this case, the glutamate residues) was divided by N_frames_, the total number of frames in the time window. The means and standard errors of the ion occupancies were calculated from non-overlapping 50 ns time windows of the 200 ns simulations. By averaging the amount of time an ion is located within the predetermined binding site, the ion residence time (tr) was calculated. The mean and standard error of tr were calculated in the same fashion as those for ion occupancies.

### Production and purification of EFL

C43 (DE3) *E. coli* cells transformed with the EFL plasmid (32) were plated onto LB agar plates supplemented with ampicillin (Amp). Plates were incubated at 37 °C for 16 h and yielded colonies. A single colony was added to a starter culture of 100 mL LB supplemented with Amp in a 250 mL conical flask before culturing for 16 h at 30 °C and 180 RPM. Hereafter, 10 mL of the starter culture was added to 1 L portions of Amp-supplemented LB in 3 L baffled flasks and incubated at 37 °C and 220 RPM. Once the cultures had grown to an OD600 of 0.6-0.8, they were induced with 1 mM IPTG. The induced culture was incubated at 37 °C and 220 RPM for 4 h before harvesting the cells by centrifugation at 6000 x *g* for 10 min at 4 °C. The resulting wet cell mass was flash-frozen in liquid N_2_ and stored at −70 °C.

The cell pellet was thawed at room temperature (20-22 °C) and resuspended in 100 mL lysis buffer (20 mM TRIS-HCl pH 8.0, 500 mM NaCl) supplemented with 50 μg lysozyme (SIGMA), and 50 μg DNase (SIGMA) per L culture. The resuspended cells were lysed using a probe sonicator (Model HD2200, Probe KE76. Bandelin) at 60 % power, 5 s on/off for 5 × 2 min rounds with 2 min rest periods in between each sonication round. Triton X-100 was added to the cell lysate to a final concentration of 2 % (v/v) before incubation with gentle rotation for 1 h at room temperature. The cell lysate was then centrifuged at 20,000 x *g* for 30 min at 4 °C. The pelleted inclusion bodies were resuspended in 100 mL TBS (20 mM TRIS-HCl pH 7.8, 140 mM NaCl) supplemented with 10 mM β-mercaptoethanol (β-ME). Empigen (SIGMA) was added to the suspension at a final concentration of 3 % (w/v). Solubilization was carried out for 18 h with stirring at room temperature. The next morning, the solubilized inclusion body material was centrifuged at 40,000 x *g* for 30 min at 15 °C and the supernatant was used for subsequent steps.

Imidazole was added to the supernatant to a final concentration of 10 mM before incubation for 1 h at room temperature with 2 mL PureCube Ni-NTA agarose resin (Cat #31105), pre-washed with 60 mL TBS supplemented with 0.3 % (w/v) Empigen, 10 mM β-ME, and 10 mM imidazole. The suspension was loaded onto a gravity column. Once all the solubilized material had passed through and the resin had settled, the bound protein was exchanged into CHAPS by washing with 50 mL 20 mM TRIS-HCl pH 7.8, 2 M NaCl, 1 % (w/v) CHAPS, 10 mM β-ME, and 10 mM imidazole. The resin was washed with TBS supplemented with 1 % (w/v) CHAPS, 10 mM β-ME and 0.04-1 M imidazole. EFL eluted during all imidazole washes with the bulk of the protein eluting at 100-500 mM imidazole. Protein eluting at 0.1 – 1 M imidazole was pooled and exchanged into TBS supplemented with 1 % (w/v) CHAPS and 1 mM DTT on a PD10 column. The sample was then concentrated in a concentrator with a 10 kDa MWCO. Concentration was determined by a combination of absorbance measurements at 280 nm (extinction coefficient = 5,960 M^−1^.cm^−1^, ProtParam) and Bradford assay. Sample homogeneity was investigated by analytical size exclusion chromatography on a Superose® 6 10/300 GL gel filtration column (GE Healthcare). Aliquots of purified EFL at 0.4 mg/mL were flash frozen in liquid nitrogen and stored at −70 °C.

EFL was purified in FC16 detergent in a similar fashion. Following cell lysis and TRITON X-100 lysate treatment as described above, the pelleted inclusion bodies were resuspended in 100 mL HBS (20 mM HEPES-NaOH pH 7.8, 500 mM NaCl) supplemented with 10 mM β-mercaptoethanol (β-ME). 1.2 g of FC16 solid (Cube Biotech (Cat #16038) was added to the suspension at a final concentration of 1.2 % (w/v). Solubilization was carried out for 18 h with stirring at room temperature. Hereafter, the solubilized inclusion body material was centrifuged at 40,000 x *g* for 30 min at 15 °C and the supernatant was used for subsequent steps. Imidazole was added to the supernatant to a final concentration of 10 mM before incubation for 1 h at room temperature with 2 mL PureCube Ni-NTA agarose resin (Cat #31105), pre-washed with 60 mL HBS supplemented with 0.05 % (w/v) FC16, 10 mM β-ME, and 10 mM imidazole. The suspension was loaded onto a gravity column. Once all the solubilized material had passed through and the resin had settled, washes with solutions containing 20 mM HEPES-NaOH pH 7.8, 2 M NaCl, 10 mM β-ME, 10 mM imidazole, and 1-0.125 % (w/v) FC16 were performed. The resin was then washed with HBS supplemented with 0.05 % (w/v) FC16, 10 mM β-ME and 0.04-1 M imidazole. EFL eluted during all imidazole washes with the bulk of the protein eluting at 100-500 mM imidazole. Protein eluting at 0.1 – 1 M imidazole was pooled and concentrated to approximately 1 mL before loading onto a S200 16/60 column equilibrated with HBS supplemented with 0.05 % (w/v) FC16 and 1 mM DTT. Fractions eluting from 68 – 78 mL were pooled and concentrated using a concentrator with a 100 kDa MWCO. Concentration was determined by a combination of absorbance measurements at 280 nm (extinction coefficient = 5,960 M^−1^.cm^−1^. ProtParam) and Bradford assay. Aliquots of purified EFL at 20-30 mg/mL were flash frozen in liquid nitrogen and stored at −70 °C.

### Mass photometry (MP)

Samples were measured on a Refeyn OneMP mass photometer with a 10.8 × 2.9 μm^2^ (128 × 35 pixels) field of view. 500 nM EFL stocks in CHAPS- or FC16-containing buffers were diluted into detergent-free buffer (50 mM HEPES pH 7.5, 150 mM NaCl, 1 mM DTT). The diluted samples containing 25-50 nM protein were then applied onto microscopy slides that had been cleaned by consecutive sonication in Milli-Q water, isopropanol, and Milli-Q water and standard mass photometry measurements (landing assays) were carried out in silicone gaskets (3 mm × 1 mm, GBL103250, Grace Bio-Labs) and images were acquired for 60 s at 331 Hz.

### Size exclusion chromatography coupled to multi-angle light scattering (SEC-MALS)

SEC-MALS experiments were performed at room temperature on an Äkta Purifier equipped with a Superdex 200, GL 10/300 column (GE Healthcare). Sample elution was monitored by using a multi-angle light scattering detector (miniDAWN TREOS, Wyatt) and a differential refractive index detector (Optilab T-rEX, Wyatt). Filtered, degassed buffer (20 mM HEPES, 500 mM NaCl, 10 mM TCEP, 0.05 % (w/v) FC16 pH 7.8) was pumped at a constant flow rate of 0.5 mL/min. The samples (300 μL) contained 0.25 mM EFL and were centrifuged at 20,000 x g at 4 °C for 15 min immediately prior to injection. The mass average molar mass across the elution peak was determined in ASTRA 6 software (Wyatt Technology) using the triple-detector method with protein dn/dc, protein UV 280 nm extinction coefficient, FC16 dn/dc, and FC16 UV 280 nm extinction coefficient set to 0.185 mL/g, 0.62 mL/mg.cm, 0.1327 mL/g, and 0.0 mL/mg.cm, respectively.

### Solid-phase synthesis of the ETM peptide

The ETM peptide, corresponding to the TM domain of SARS-CoV-2 (residues 8-38), was produced by standard solid phase FMOC chemistry on a Liberty Blue HT12 automated microwave peptide synthesizer. The peptide was cleaved from the resin by washing with a mixture 95/ 2.5/2.5 % (by vol.) mixture of trifluoroacetic acid (TFA), N-tris(hydroxymethyl)methyl-2-aminoethanesulfonic acid (TES), and water over 2.5 h. Residual salts remaining after cleavage were removed by repeated suspension of the water-insoluble peptide solid in water. The peptide was pelleted by centrifugation and lyophilized overnight. ETM mass analysis, performed with an Agilent 6460 Triple Quadrupole LC/MS, confirmed the predicted peptide molecular weight to within 1 Da (Fig. S3A). ETM was reconstituted into TBS supplemented with 1 %(w/v) CHAPS and 1 mM DTT to 0.4 mg/mL by sonication for 30 min at 30 °C.

### Planar Lipid Bilayer Formation and incorporation of SARS-CoV-2 E constructs

EFL and ETM were incorporated into bovine phosphatidylethanolamine (PE) (Avanti Polar Lipids, USA) lipid bilayers under voltage-clamp conditions using previously described techniques (71). Briefly, planar lipid bilayers were formed across a 150 μm diameter aperture in a Delrin cuvette that separates the *cis* and the *trans* 1 mL compartments (see Fig. 1B for schematic). Protein samples were diluted 1:1 in 400 mM sucrose, 5 mM HEPES, pH 7.2 with Tris-(hydroxymethyl) aminoethane (VWR Chemicals, Leuven Belgium; cat no. 103156X), hereafter known as TRIS and approximately 0.15 - 0.3 μg (w) was added to the *cis* chamber. Incorporation was facilitated by constant stirring. Protein incorporation was assessed by current fluctuations away from 0 mV. Following incorporation, solution in the *cis* chamber was exchanged 10 times to remove excess protein. The *trans* chamber was held at ground, and the *cis* chamber was voltage-clamped at various potentials relative to ground. Bilayers with capacitances between 70 and 115 pF were used in all experiments and stability was checked periodically throughout the experiment. The transmembrane current was recorded under voltage-clamp conditions using a BC-525C amplifier (Warner Instruments, Harvard). Channel recordings were low-pass filtered at 10 kHz with a four-pole Bessel filter, digitized at 100 kHz using a National Instruments acquisition interface (NIDAQ-MX, National Instruments, Austin, TX) and recorded on a computer hard drive using WinEDR software version 4.0.0 or later (John Dempster, University of Strathclyde, UK). Current fluctuations were recorded over time and subsequently filtered at either 200 or 800 Hz (−3 dB) using the low pass digital filter within WinEDR. All single channel analysis was carried out using WinEDR.

Where multiple channels had been incorporated into the bilayer the mean current was calculated using noise analysis. 800 Hz low pass filtered recordings were utilized to find the mean current by plotting current fluctuations subdivided into *N* samples over time with the following equation:

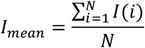

Where *I(i*) is the *i^th^* of *N* samples within the recording. Mean direct current was taken from recordings ≥30 s long and the calculated average was then plotted as a function of voltage.

Predicted reversal potentials (E_rev_) of a given ion were calculated using the Nernst equation. The relative permeability of monovalent cations was calculated using the Goldman-Hodgkin-Katz equation (72–74):

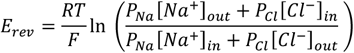

where R is the ideal gas constant (8.314 Jmol^−1^), T is the temperature in Kelvin (293.15 K), and F is the Faraday constant (9.6485 × 10^4^ C mol^−1^). The relative Ca^2+^ to Na^+^ permeability ratio (PCa^2+^/PNa^+^) was calculated using a previously described modified Fatt-Ginsborg equation (74–77):

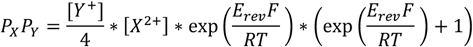

where R, T, and F have their usual values and meanings and E_rev_ is the experimentally determined reversal potential. X refers to ions in the *trans* chamber and Y refers to ions in the *cis* chamber. [X] and [Y] refer to the concentrations of the ions. (E_rev_) is the zero current reversal potential. The reversal potential was taken to be the voltage at which no current fluctuations were observed. Junction potentials were calculated using Clampex software version 10.2 (Molecular Devices) and subtracted from the reversal potential obtained from each experiment.

BioRender.com was utilized for schematics and infographics and is cited in figure legends where it was used. All statistics and quantitative figures were run/generated in GraphPad Prism 9.0 or later. A one-way non-parametric ANOVA on ranks (Kruskal-Wallis test), with Dunn’s multiple comparisons post-tests were run. Significance was determined if p<0.05. All error bars and the ± symbol indicate standard deviation of the mean (sd).

### *In vitro* solutions

For experiments where Na^+^ is the permeant ion, the *trans* chamber contained 210 mM NaCl, 10 mM HEPES pH 7.2 and the *cis* chamber contained 410 mM NaCl, 10 mM HEPES pH 7.2. For Ca^2+^ permeation studies, the *trans* chamber contained 210 mM CaCl_2_, 10 mM HEPES pH 7.2 and the *cis* chamber contained 510 mM CaCl_2_, 10 mM HEPES pH 7.2. For Na^+^:Ca^2+^ permeability ratio experiments the *cis* and *trans* chambers were perfused resulting in a concentration of 210 mM Na^+^ in the *cis* chamber and 210 mM Ca^2+^ in the *trans* chamber.

The pH of all solutions was measured at room temperature (20 °C) using a Hannah pH 211 microprocessor pH meter and pH electrode (Hannah Instruments Ltd., UK). The free [Ca^2+^] of all Na^+^ solutions was determined using a Ca^2+^ electrode. An HI 4004 Calcium Half-Cell (HANNA, RI, USA) and HI 4104 Calcium Combination Electrode (HANNA, RI, USA) were used on a HANNA pH 211 Microprocessor pH Meter as described previously (69, 70). The free Ca^2+^ concentration of the 210 mM NaCl, 10 mM HEPES, pH 7.2 solution was 3.873 ± 0.711 (sd) μM (*n* = 4). The 1 M NaCl solution which was used to create the asymmetric gradient in the *cis* chamber was 21.043 ± 0.463 (sd) μM (*n* = 3). The *cis* chamber contained approximately 6.791 μM free Ca^2+^ on average when Na^+^ was the permeant ion and the *trans* chamber contained 3.873 μM free Ca^2+^ when Na^+^ was the permeant ion.

All chemicals were AnalaR or the best equivalent grade from BDH (Poole, UK) or SigmaAldrich (Dorset, UK). All solutions were made in milliQ ultrapure de-ionized water and those for use in bilayer experiments were filtered through a Millipore membrane filter (0.45-μm pore). Any chemicals or reagents originating from other sources are indicated in the methods

#### Ca^2+^ activation experiments

To examine the effect of luminal Ca^2+^ on the gating characteristics of the SARS-CoV-2 E protein, the EFL channel was incorporated into the bilayer and Ca^2+^ was sequentially added at concentrations of 0, 50, 100, 300, 700, 1000 μM to the *trans* chamber in the presence of a Na^+^ gradient. Changes to gating characteristics were monitored by recording the channel for **≥** 60 s at each *trans* Ca^2+^ concentration. The bilayer was voltage-clamped at +60 mV (approximately +63.1 mV with junction potential correction assuming Na^+^ is the permeant ion) and Na^+^ was the permeant ion (410 mM NaCl *cis*: 210 mM NaCl *trans*). To determine the % current increase in picoamperes (pA), the following formula was applied to normalize the raw continuous recordings: (B - A)/A, where B is the experimental condition and A is the control condition.

#### Elution buffer controls

Elution buffer contained 20 mM TRIS-HCl, 140 mM NaCl, 1 % (w/v) CHAPS, and 0.25 mM DTT, pH 7.5. Elution buffer was diluted 1:1 with a sucrose solution. The sucrose solution contained 400 mM sucrose, 5 mM HEPES with pH adjusted to 7.2 with TRIS. Following formation of a bilayer in 210 mM NaCl *trans*: 410 NaCl *cis* solutions an equivalent volume of elution buffer used in our protein incorporation studies was added to the *cis* chamber. The *cis* chamber was stirred for a minimum of 10-15 mins while the membrane voltage was clamped between +120 or −120 mV after which a voltage ramp was applied by alternating the holding command from −60 to +60 mV (−56.9 mV to +63.1 mV with junction potential correction imposed) in 10 mV steps. Each voltage was held for ≥ 20 seconds.

## Acknowledgements

We thank the University of Dundee I.T. services for maintenance of the School of Life Sciences high-performance computing (HPC) cluster which is utilized in this research. Furthermore, we would like to thank Neil J. Thomson for assistance with coding and figure creation. The project was supported by the UKRI Biotechnology and Biological Sciences Research Council (BBSRC; grant number BB/V01997X/1). QWH was further supported by the Wellcome Trust Institutional Strategic Support Funds (grant number 204821/Z/16/Z).

## Contributions

Research study conceptualization and design: MC, SJP, UZ. Computational experiments: LHA, CMI. Laboratory experiments/analysis: KR, QWH, NO, ES. Data analysis: LHA, KR, QWH, CMI. Supervision: MC, SJP, UZ. Writing/editing of the manuscript: LHA, KR, QWH, MC, SJP, UZ.

## Supplementary Information

Figures S1-S10, Tables S1-S6, and an atomistic model of fully-palmitoylated, ion-conductive SARS-CoV-2 E with a view to informing anti-viral drug design efforts.

## References

1. Callaway E. What Omicron’s BA. 4 and BA. 5 variants mean for the pandemic. Nature. 2022;606:848–9. https://doi.org/10.1038/d41586-022-01730-y.

2. Araf Y, Akter F, Tang Y-d, et al. Omicron variant of SARS-CoV-2: Genomics, transmissibility, and responses to current COVID-19 vaccines. Journal of Medical Virology. 2022;94(5):1825–32. https://doi.org/10.1002/jmv.27588.

3. Tian D, Sun Y, Xu H, Ye Q. The emergence and epidemic characteristics of the highly mutated SARS-CoV-2 Omicron variant. Journal of Medical Virology. 2022;94(6):2376–83. https://doi.org/10.1002/jmv.27643.

4. Zhang X, Wu S, Wu B, et al. SARS-CoV-2 Omicron strain exhibits potent capabilities for immune evasion and viral entrance. Signal Transduction and Targeted Therapy. 2021;6(1):430. https://www.doi.org/10.1038/s41392-021-00852-5.

5. Masters PS. The molecular biology of coronaviruses. Advances in virus research. 2006;66:193–292. https://doi.org/10.1016/S0065-3527(06)66005-3.

6. Jiang S, Du L, Shi Z. An emerging coronavirus causing pneumonia outbreak in Wuhan, China: calling for developing therapeutic and prophylactic strategies. Emerging Microbes & Infections. 2020;9(1):275–7. https://www.doi.org/10.1080/22221751.2020.1723441.

7. Xia X. Domains and Functions of Spike Protein in SARS-Cov-2 in the Context of Vaccine Design. Viruses. 2021;13(1):109. https://www.doi.org/10.3390/v13010109.

8. Huang Y, Yang C, Xu X-f, Xu W, Liu S-w. Structural and functional properties of SARS-CoV-2 spike protein: potential antivirus drug development for COVID-19. Acta Pharmacologica Sinica. 2020;41(9):1141–9. https://www.doi.org/10.1038/s41401-020-0485-4.

9. Wen W, Chen C, Tang J, et al. Efficacy and safety of three new oral antiviral treatment (molnupiravir, fluvoxamine and Paxlovid) for COVID-19: a meta-analysis. Annals of Medicine. 2022;54(1):516–23. https://www.doi.org/10.1080/07853890.2022.2034936.

10. Hsu J-N, Chen J-S, Lin S-M, et al. Targeting the N-Terminus Domain of the Coronavirus Nucleocapsid Protein Induces Abnormal Oligomerization via Allosteric Modulation. Frontiers in Molecular Biosciences. 2022;9. https://www.doi.org/10.3389/fmolb.2022.871499.

11. Park SH, Siddiqi H, Castro DV, et al. Interactions of SARS-CoV-2 envelope protein with amilorides correlate with antiviral activity. PLOS Pathogens. 2021;17(5):e1009519. https://doi.org/10.1371/journal.ppat.1009519.

12. Schoeman D, Fielding BC. Coronavirus envelope protein: current knowledge. Virology Journal. 2019;16(1):69. https://www.doi.org/10.1186/s12985-019-1182-0.

13. Dey D, Borkotoky S, Banerjee M. In silico identification of Tretinoin as a SARS-CoV-2 envelope (E) protein ion channel inhibitor. Comput Biol Med. 2020;127:104063-. https://www.doi.org/10.1016/j.compbiomed.2020.104063.

14. Cao Y, Yang R, Wang W, et al. Computational Study of the Ion and Water Permeation and Transport Mechanisms of the SARS-CoV-2 Pentameric E Protein Channel. Frontiers in Molecular Biosciences. 2020;7(270). https://www.doi.org/10.3389/fmolb.2020.565797.

15. Gupta MK, Vemula S, Donde R, Gouda G, Behera L, Vadde R. In-silico approaches to detect inhibitors of the human severe acute respiratory syndrome coronavirus envelope protein ion channel. J Biomol Struct Dyn. 2021;39(7):2617–27. https://www.doi.org/10.1080/07391102.2020.1751300.

16. Sun S, Karki C, Aguilera J, Lopez Hernandez AE, Sun J, Li L. Computational Study on the Function of Palmitoylation on the Envelope Protein in SARS-CoV-2. Journal of Chemical Theory and Computation. 2021;17(10):6483–90. https://www.doi.org/10.1021/acs.jctc.1c00359.

17. Mehregan A, Pérez-Conesa S, Zhuang Y, et al. Probing effects of the SARS-CoV-2 E protein on membrane curvature and intracellular calcium. Biochim Biophys Acta Biomembr. 2022;1864(10):183994. https://www.doi.org/10.1016/j.bbamem.2022.183994.

18. Ren SY, Wang WB, Gao RD, Zhou AM. Omicron variant (B.1.1.529) of SARS-CoV-2: Mutation, infectivity, transmission, and vaccine resistance. World J Clin Cases. 2022;10(1):1–11. https://www.doi.org/10.12998/wjcc.v10.i1.1.

19. Gupta RK, Topol EJ. COVID-19 vaccine breakthrough infections. Science. 2021;374(6575):1561–2. https://www.doi.org/10.1126/science.abl8487.

20. Wang R, Chen J, Gao K, Wei G-W. Vaccine-escape and fast-growing mutations in the United Kingdom, the United States, Singapore, Spain, India, and other COVID-19-devastated countries. Genomics. 2021;113(4):2158–70. https://doi.org/10.1016/j.ygeno.2021.05.006.

21. Wang C, Liu Z, Chen Z, et al. The establishment of reference sequence for SARS-CoV-2 and variation analysis. Journal of Medical Virology. 2020;92(6):667–74. https://doi.org/10.1002/jmv.25762.

22. Xia B, Shen X, He Y, et al. SARS-CoV-2 envelope protein causes acute respiratory distress syndrome (ARDS)-like pathological damages and constitutes an antiviral target. Cell Research. 2021;31(8):847–60. https://www.doi.org/10.1038/s41422-021-00519-4.

23. Verdiá-Báguena C, Nieto-Torres JL, Alcaraz A, et al. Coronavirus E protein forms ion channels with functionally and structurally-involved membrane lipids. Virology. 2012;432(2):485–94. https://www.doi.org/10.1016/j.virol.2012.07.005.

24. Pervushin K, Tan E, Parthasarathy K, et al. Structure and Inhibition of the SARS Coronavirus Envelope Protein Ion Channel. PLOS Pathogens. 2009;5(7):e1000511. https://www.doi.org/10.1371/journal.ppat.1000511.

25. Nieto-Torres JL, Verdiá-Báguena C, Jimenez-Guardeño JM, et al. Severe acute respiratory syndrome coronavirus E protein transports calcium ions and activates the NLRP3 inflammasome. Virology. 2015;485:330–9. https://www.doi.org/10.1016/j.virol.2015.08.010.

26. Wilson L, McKinlay C, Gage P, Ewart G. SARS coronavirus E protein forms cation-selective ion channels. Virology. 2004;330(1):322–31. https://doi.org/10.1016/j.virol.2004.09.033.

27. DeDiego ML, Nieto-Torres JL, Regla-Nava JA, et al. Inhibition of NF-kB-Mediated Inflammation in Severe Acute Respiratory Syndrome Coronavirus-Infected Mice Increases Survival. Journal of Virology. 2014;88(2):913–24. https://www.doi.org/10.1128/JVI.02576-13.

28. Jimenez-Guardeño JM, Nieto-Torres JL, DeDiego ML, et al. The PDZ-Binding Motif of Severe Acute Respiratory Syndrome Coronavirus Envelope Protein Is a Determinant of Viral Pathogenesis. PLOS Pathogens. 2014;10(8):e1004320. https://www.doi.org/10.1371/journal.ppat.1004320.

29. Verdiá-Báguena C, Aguilella VM, Queralt-Martín M, Alcaraz A. Transport mechanisms of SARS-CoV-E viroporin in calcium solutions: Lipid-dependent Anomalous Mole Fraction Effect and regulation of pore conductance. Biochim Biophys Acta Biomembr. 2021;1863(6):183590. https://www.doi.org/10.1016/j.bbamem.2021.183590.

30. Singh Tomar PP, Arkin IT. SARS-CoV-2 E protein is a potential ion channel that can be inhibited by Gliclazide and Memantine. Biochem Biophys Res Commun. 2020;530(1):10–4. https://www.doi.org/10.1016/j.bbrc.2020.05.206.

31. Cabrera-Garcia D, Bekdash R, Abbott GW, Yazawa M, Harrison NL. The envelope protein of SARS-CoV-2 increases intra-Golgi pH and forms a cation channel that is regulated by pH. The Journal of Physiology. 2021;599(11):2851–68. https://doi.org/10.1113/JP281037.

32. Hutchison JM, Capone R, Luu DD, et al. Recombinant SARS-CoV-2 envelope protein traffics to the <em>trans</em>-Golgi network following amphipol-mediated delivery into human cells. Journal of Biological Chemistry. 2021;297(2). https://www.doi.org/10.1016/j.jbc.2021.100940.

33. Young G, Hundt N, Cole D, et al. Quantitative mass imaging of single biological macromolecules. Science. 2018;360(6387):423–7. https://www.doi.org/10.1126/science.aar5839.

34. Foley EDB, Kushwah MS, Young G, Kukura P. Mass photometry enables label-free tracking and mass measurement of single proteins on lipid bilayers. Nature Methods. 2021;18(10):1247–52. https://www.doi.org/10.1038/s41592-021-01261-w.

35. Gimpl K, Klement J, Keller S. Characterising protein/detergent complexes by triple-detection size-exclusion chromatography. Biological Procedures Online. 2016;18(1):4. https://www.doi.org/10.1186/s12575-015-0031-9.

36. Toft-Bertelsen TL, Jeppesen MG, Tzortzini E, et al. Amantadine inhibits known and novel ion channels encoded by SARS-CoV-2 in vitro. Communications Biology. 2021;4(1):1347. https://doi.org/10.1038/s42003-021-02866-9.

37. Surya W, Li Y, Torres J. Structural model of the SARS coronavirus E channel in LMPG micelles. Biochimica et Biophysica Acta (BBA) - Biomembranes. 2018;1860(6):1309–17. https://doi.org/10.1016/j.bbamem.2018.02.017.

38. Mehregan A, Pérez-Conesa S, Zhuang Y, et al. Probing effects of the SARS-CoV-2 E protein on membrane curvature and intracellular calcium. Biochimica et Biophysica Acta (BBA)-Biomembranes. 2022;1864(10):183994. https://doi.org/10.1016/j.bbamem.2022.183994.

39. Aryal P, Sansom MS, Tucker SJ. Hydrophobic gating in ion channels. Journal of molecular biology. 2015;427(1):121–30. https://doi.org/10.1016/j.jmb.2014.07.030.

40. Beckstein O, Biggin PC, Sansom MS. A hydrophobic gating mechanism for nanopores. The Journal of Physical Chemistry B. 2001;105(51):12902–5. https://doi.org/10.1021/jp012233y.

41. Medeiros-Silva J, Somberg NH, Wang HK, et al. pH-and Calcium-Dependent Aromatic Network in the SARS-CoV-2 Envelope Protein. Journal of the American Chemical Society. 2022;144(15):6839–50. https://www.doi.org/10.1021/jacs.2c00973.

42. Fung TS, Liu DX. Post-translational modifications of coronavirus proteins: roles and function. Future Virology. 2018;13(6):405 – 30. https://www.doi.org/10.2217/fvl-2018-0008.

43. Lopez LA, Riffle AJ, Pike SL, Gardner D, Hogue BG. Importance of Conserved Cysteine Residues in the Coronavirus Envelope Protein. Journal of Virology. 2008;82(6):3000–10. www.doi.org/10.1128/JVI.01914-07.

44. Sobocińska J, Roszczenko-Jasińska P, Ciesielska A, Kwiatkowska K. Protein palmitoylation and its role in bacterial and viral infections. Frontiers in immunology. 2018;8:2003. https://doi.org/10.3389/fimmu.2017.02003.

45. Blaskovic S, Blanc M, van der Goot FG. What does S-palmitoylation do to membrane proteins? The FEBS Journal. 2013;280(12):2766–74. https://doi.org/10.1111/febs.12263.

46. Paroutis P, Touret N, Grinstein S. The pH of the Secretory Pathway: Measurement, Determinants, and Regulation. Physiology. 2004;19(4):207–15. https://www.doi.org/10.1152/physiol.00005.2004.

47. Wang X, Kirberger M, Qiu F, Chen G, Yang JJ. Towards predicting Ca2+-binding sites with different coordination numbers in proteins with atomic resolution. Proteins. 2009;75(4):787–98. https://www.doi.org/10.1002/prot.22285.

48. Mandala VS, McKay MJ, Shcherbakov AA, Dregni AJ, Kolocouris A, Hong M. Structure and drug binding of the SARS-CoV-2 envelope protein transmembrane domain in lipid bilayers. Nature Structural & Molecular Biology. 2020;27(12):1202–8.https://www.doi.org/10.1038/s41594-020-00536-8.

49. Torres J, Maheswari U, Parthasarathy K, Ng L, Liu DX, Gong X. Conductance and amantadine binding of a pore formed by a lysine-flanked transmembrane domain of SARS coronavirus envelope protein. Protein Sci. 2009;16(9):2065–71. https://www.doi.org/10.1110/ps.062730007.

50. Harrison NL, Abbott GW, Gentzsch M, et al. How many SARS-CoV-2 “viroporins” are really ion channels? Communications Biology. 2022;5(1):859. https://www.doi.org/10.1038/s42003-022-03669-2.

51. Appenzeller-Herzog C, Hauri H-P. The ER-Golgi intermediate compartment (ERGIC): in search of its identity and function. Journal of Cell Science. 2006;119(11):2173–83. https://www.doi.org/10.1242/jcs.03019.

52. Zhang L-k, Sun Y, Zeng H, et al. Calcium channel blocker amlodipine besylate is associated with reduced case fatality rate of COVID-19 patients with hypertension. medRxiv. 2020. https://www.doi.org/10.1101/2020.04.08.20047134.

53. Solaimanzadeh I. Nifedipine and Amlodipine Are Associated With Improved Mortality and Decreased Risk for Intubation and Mechanical Ventilation in Elderly Patients Hospitalized for COVID-19. Cureus. 2020;12(5):e8069–e. https://www.doi.org/10.7759/cureus.8069.

54. Choksi TT, Zhang H, Chen T, Malhotra N. Outcomes of Hospitalized COVID-19 Patients Receiving Renin Angiotensin System Blockers and Calcium Channel Blockers. American Journal of Nephrology. 2021;52(3):250–60. https://www.doi/org/10.1159/000515232.

55. Yan F, Huang F, Xu J, et al. Antihypertensive drugs are associated with reduced fatal outcomes and improved clinical characteristics in elderly COVID-19 patients. Cell Discovery. 2020;6(1):77. https://doi.org/10.1038/s41421-020-00221-6.

56. Berlansky S, Sallinger M, Grabmayr H, et al. Calcium Signals during SARS-CoV-2 Infection: Assessing the Potential of Emerging Therapies. Cells. 2022;11(2). https://www.doi.org/10.3390/cells11020253.

57. Lee J, Cheng X, Swails JM, et al. CHARMM-GUI Input Generator for NAMD, GROMACS, AMBER, OpenMM, and CHARMM/OpenMM Simulations Using the CHARMM36 Additive Force Field. Journal of Chemical Theory and Computation. 2016;12:405–13. https://www.doi.org/10.1021/acs.jctc.5b00935.

58. Van Der Spoel D, Lindahl E, Hess B, Groenhof G, Mark AE, Berendsen HJ. GROMACS: fast, flexible, and free. J Comput Chem. 2005;26(16):1701–18. https://www.doi.org/10.1002/jcc.20291.

59. Zhang A, Yu H, Liu C, Song C. The Ca2+ permeation mechanism of the ryanodine receptor revealed by a multi-site ion model. Nature Communications. 2020;11(1):922. https://www.doi.org/10.1038/s41467-020-14573-w.

60. Evans DJ, Holian BL. The Nose–Hoover thermostat. The Journal of Chemical Physics. 1985;83(8):4069–74. https://www.doi.org/10.1063/1.449071.

61. Parrinello M, Rahman A. Polymorphic transitions in single crystals: A new molecular dynamics method. Journal of Applied Physics. 1981;52(12):7182–90. https://www.doi.org/10.1063/1.328693.

62. Darden T, York D, Pedersen L. Particle mesh Ewald: An N·log(N) method for Ewald sums in large systems. The Journal of Chemical Physics. 1993;98(12):10089–92. https://www.doi.org/10.1063/1.464397.

63. Hess B, Bekker H, Berendsen HJC, Fraaije JGEM. LINCS: A linear constraint solver for molecular simulations. Journal of Computational Chemistry. 1997;18(12):1463–72. https://doi.org/10.1002/(SICI)1096-987X(199709)18:12<1463::AID-JCC4>3.0.CO;2-H.

64. Gumbart J, Khalili-Araghi F, Sotomayor M, Roux B. Constant electric field simulations of the membrane potential illustrated with simple systems. Biochimica et Biophysica Acta (BBA) - Biomembranes. 2012;1818(2):294–302. https://doi.org/10.1016/j.bbamem.2011.09.030.

65. Aksimentiev A, Schulten K. Imaging α-Hemolysin with Molecular Dynamics: Ionic Conductance, Osmotic Permeability, and the Electrostatic Potential Map. Biophys J. 2005;88(6):3745–61. https://doi.org/10.1529/biophysj.104.058727.

66. Gowers RJ, Linke M, Barnoud J, et al. MDAnalysis: A Python Package for the Rapid Analysis of Molecular Dynamics Simulations. Proceedings of the 15th Python in Science Conf (SCIPY2016). 2016:98–105. https://www.doi.org/10.25080/majora-629e541a-00e.

67. Klesse G, Rao S, Sansom MSP, Tucker SJ. CHAP: A Versatile Tool for the Structural and Functional Annotation of Ion Channel Pores. J Mol Biol. 2019;431 (17):3353–65. https://www.doi.org/10.1016/j.jmb.2019.06.003.

68. Hunter JD. Matplotlib: A 2D Graphics Environment. Computing in Science & Engineering. 2007;9(3):90–5.https://www.doi.org/10.1109/MCSE.2007.55.

69. Humphrey W, Dalke A, Schulten K. VMD: visual molecular dynamics. J Mol Graph. 1996;14(1):33–8, 27–8. https://www.doi.org/10.1016/0263-7855(96)00018-5.

70. Vögele M, Thomson N, Truong S, McAvity J. PENSA. Zenodo. 2021. http://doi.org/10.5281/zenodo.4362136.

71. Mueller P, Rudin DO, Ti Tien H, Wescott WC. Reconstitution of Cell Membrane Structure in vitro and its Transformation into an Excitable System. Nature. 1962;194(4832):979–80. https://www.doi.org/10.1038/194979a0.

72. Goldman DE. Potential, impedance, and rectification in membranes. THe Journal of General Physiology. 1943:37–60. https://rupress.org/jgp/article-pdf/27/1/37/1239495/37.pdf.

73. Hodgkin AL, Katz B. The Effect of Sodium Ions on the Electrical Activity of the Giant Axon of the Squid. Journal of Physiology. 1949;108:37–77. https://doi.org/10.1113/jphysiol.1949.sp004310

74. Hille B. Ion channels of excitable membranes (Sinauer, Sunderland, MA)2001. https://doi.org/10.1002/jnr.490130415.

75. Fatt P, Ginsborg BL. The ionic requirements for the production of action potentials in crustacean muscle fibres. The Journal of Physiology. 1958;142(3):516–43. https://doi.org/10.1113/jphysiol.1958.sp006034.

76. Tinker A, Williams AJ. Divalent cation conduction in the ryanodine receptor channel of sheep cardiac muscle sarcoplasmic reticulum. Journal of General Physiology. 1992;100(3):479–93. https://www.doi.org/10.1085/jgp.100.3.479.

77. Attwell D, Jack J. The interpretation of membrane current-voltage relations: A Nernst-Planck analysis. Progress in Biophysics and Molecular Biology. 1979;34:81–107. https://doi.org/10.1016/0079-6107(79)90015-4.

