## Supplemental Information for "The SARS-CoV-2 envelope (E) protein forms a calcium- and voltage-activated calcium channel"

1 10 20 30

MYSFVSEETG TLIVNSVLLF LAFVVFLLVT LAILTALRLC

40 50 60 70

AYCCNIVNVS LVKPSFYVYS RVKNLNSSRV PDLLV

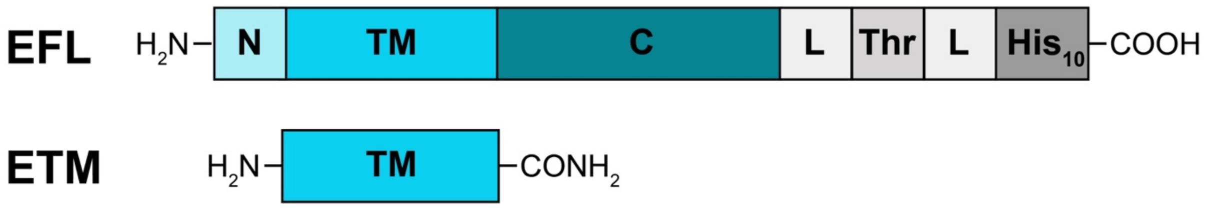

Figure S1:Amino acid sequence of the SARS-CoV-2 E protein (upper) and schematics of the E protein constructs used in this work (lower). N-terminal (N), transmembrane (TM), and C-terminal (C) domains are indicated, along with linkers (L), thrombin cleavage sites (Thr), and His10 tags (His10).

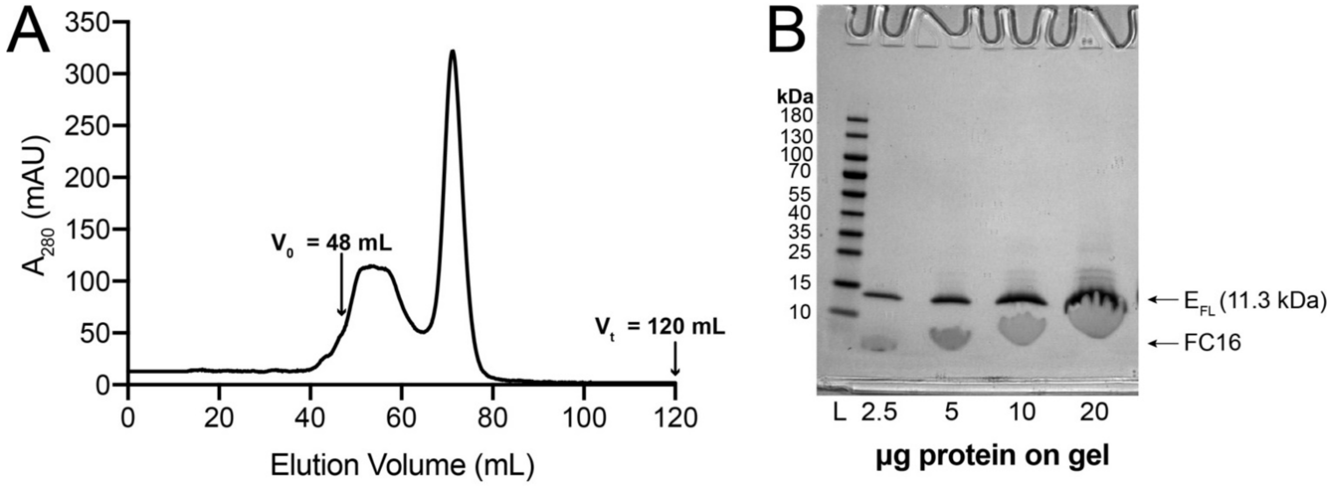

Figure S2: (A) Preparative gel filtration profile for EFL purified in 20 mM HEPES-NaOH pH 7.8, 500 mM NaCl, 0.05 %(w/v) FC16, 1 mM DTT. Gel filtration column (S200 16/60, GE Healthcare) void ($V_0$) and total ($V_t$) volumes are indicated. (B) SDS PAGE (Coomassie stain) loading series for pooled and concentrated SEC fractions eluting at 68 – 78 mL.

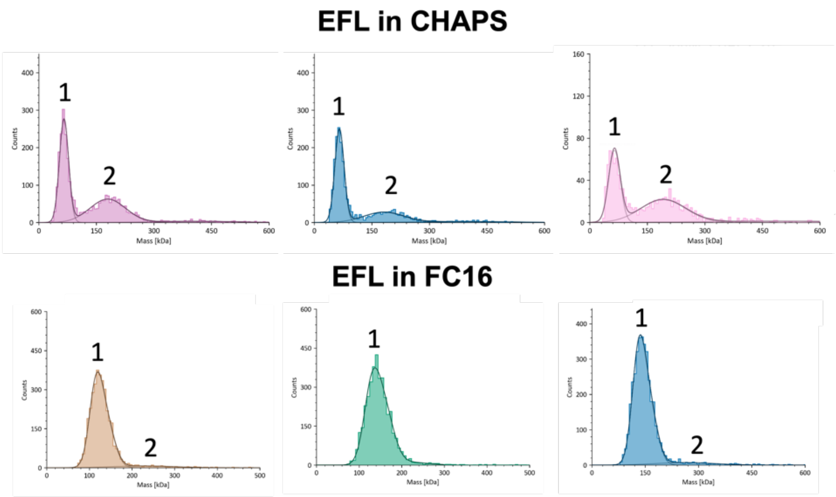

| Replicate | Estimated MWt (kDa) 1 | Estimated MWt (kDa) 2 |
|---|---|---|
| **EFL in CHAPS** | | |
| 1 | 66 ± 11.9 | 179 ± 46 |
| 2 | 65 ± 12.1 | 181 ± 52 |
| 3 | 64 ± 15.0 | 191 ± 59 |
| **EFL in FC16** | | |
| 1 | 124 ± 21 | 227 ± 69 |
| 2 | 143 ± 31 | - |
| 3 | 141 ± 25 | 263 ± 82 |

Figure S3: Mass photometry profiles for samples of EFL in CHAPS or FC16 micelles. Protein concentration was 25-50 nM.

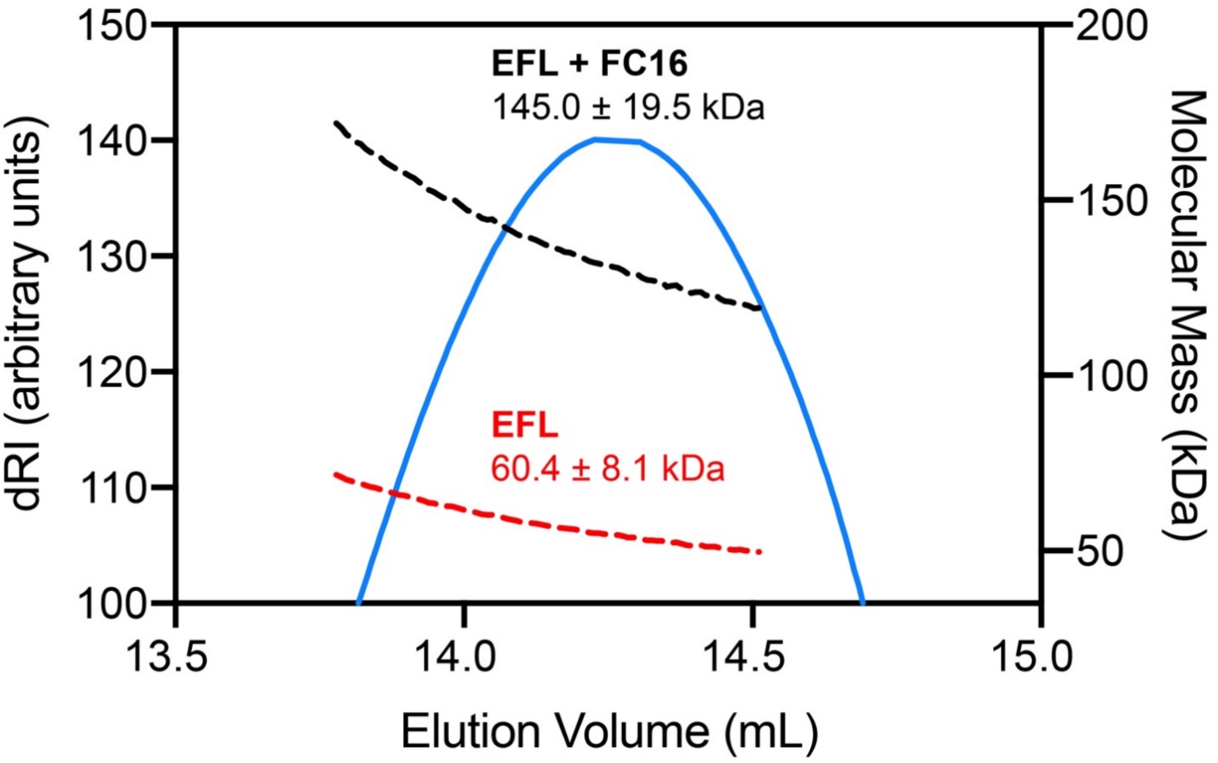

Figure S4: SEC-MALS analysis of EFL in FC16 micelles with dashed and solid lines corresponding to mass average molar masses and differential refractive index (dRI) elution profiles, respectively.

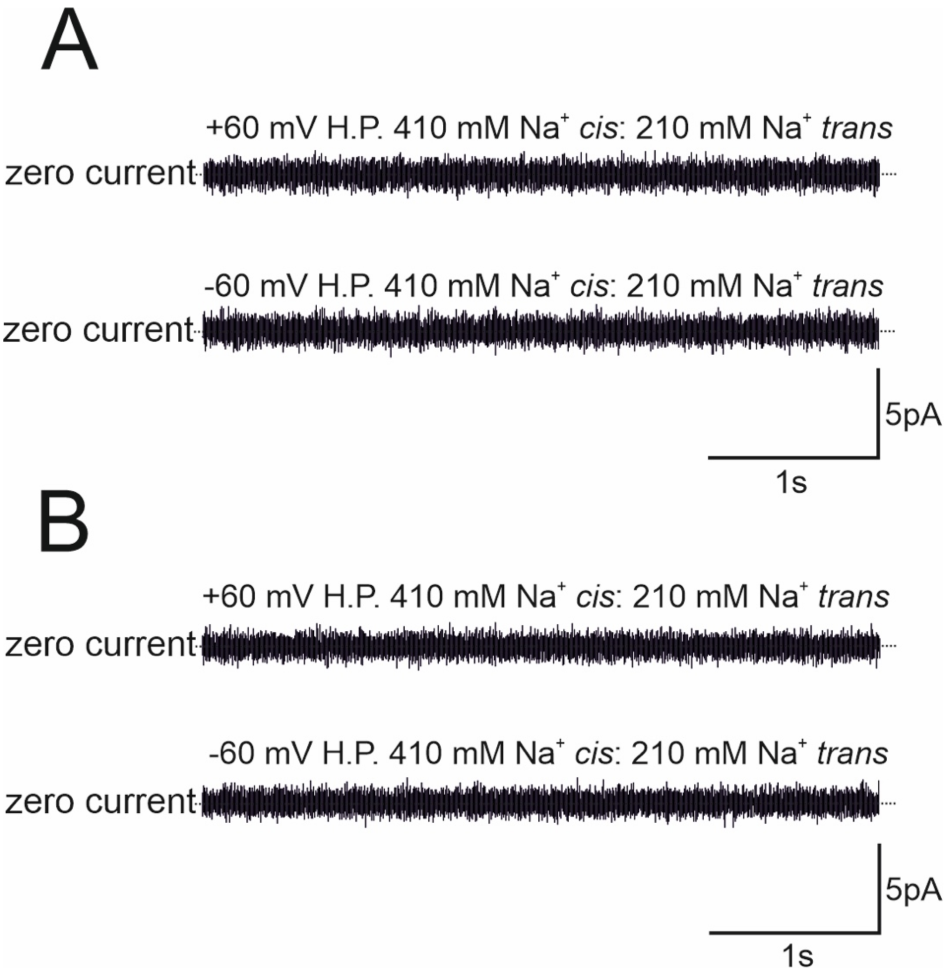

Figure S5: Control voltage clamp measurements made on planar lipid bilayer with elution buffer. (A) Continuous recording of an empty bilayer at +60 and – 60 mV. Experimental conditions are as indicated in the figure. (B) Continuous recordings of attempted Etm incorporation into the same bilayer as shown in A. Incorporation was attempted for ≥10 min with continuous stirring. Extreme voltage pulses and holding potentials were applied to thoroughly expose the bilayer to the elution buffer components. Five independent experimental repeats were conducted and zero current was observed for each experiment. Conditions are as indicated in the figure.

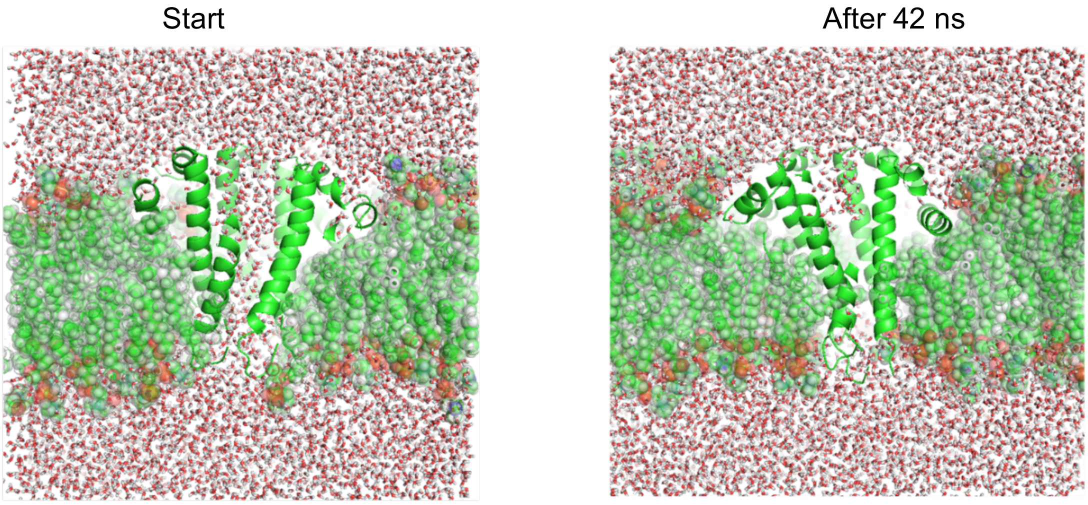

Figure S6: Dewetting of the E channel around the hydrophobic motif.

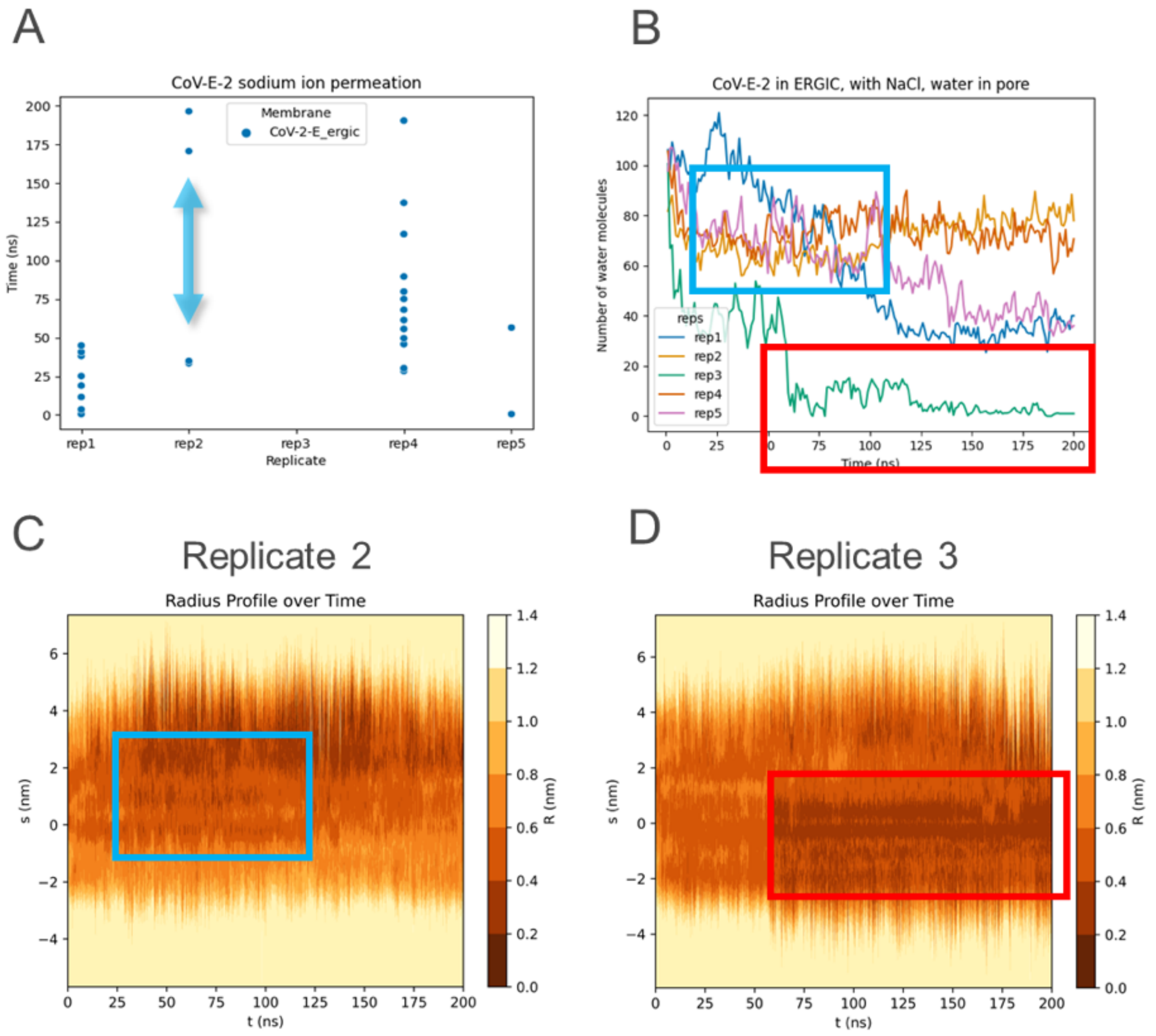

Figure S7: Dewetting and rewetting transitions in simulations with NaCl. (A) Permeation events of $Na^+$ through the E protein embedded in ERGIC-mimetic membranes Five replicates of each setup were conducted. The cyan arrow indicates the time period in replicate simulation 2 where the pore radius became too narrow to facilitate ion permeation. **(B)** Graph showing the number of water molecules occupying the transmembrane region of the pore. The cyan rectangle highlights the decrease in water molecules in replicate 2, which is caused by a decrease in pore radius. The red rectangle indicates the decrease in water when the channel is completely closed and dewetted. **(C)** Plot showing the radius profile of the pore in replicate simulation 2 throughout the simulation. Darker colors denote a smaller pore radius. The cyan rectangle shows the decrease in pore radius which causes the decrease in pore water content shown in (B) and the halt of permeation shown in (A). (D) **)** Plot showing the radius profile of the pore in replicate simulation 3 throughout the simulation. The red rectangle shows the decrease in pore radius which is caused by the decrease in pore water content shown in (B)

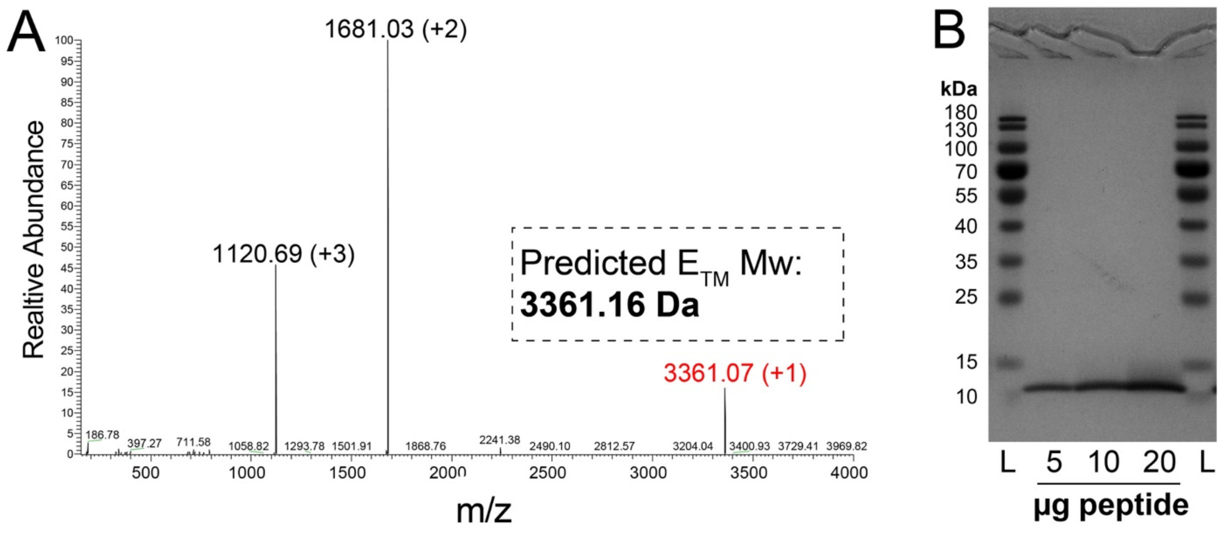

Figure S8: (A) ESI+ mass spectrum of the ETM peptide. (B) Coomassie-stained SDS PAGE loading series of ETM solubilized in CHAPS buffer (20 mM TRIS-HCl pH 7.8, 140 mM NaCl, 1 %(w/v)CHAPS, 1 mM DTT).

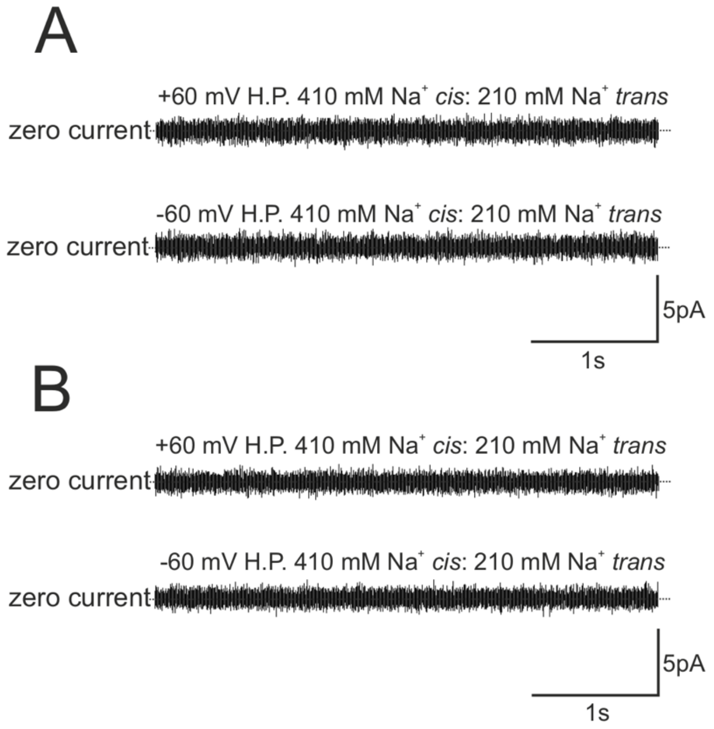

Figure S9 The SARS-Cov2-E Transmembrane domain (ETM) is non-functional. (A) Continuous recording of an empty bilayer at +60 and – 60 mV (with junction potential correction imposed –56.9 mV to +63.1 mV). Experimental conditions are as indicated in the figure. (B) Continuous recordings of attempted ETM incorporation into the same bilayer as shown in A. Incorporation was attempted for ≥20 min with continuous stirring. Seven independent experimental repeats were conducted and zero current was observed for each experiment. Conditions are as indicated in the figure.

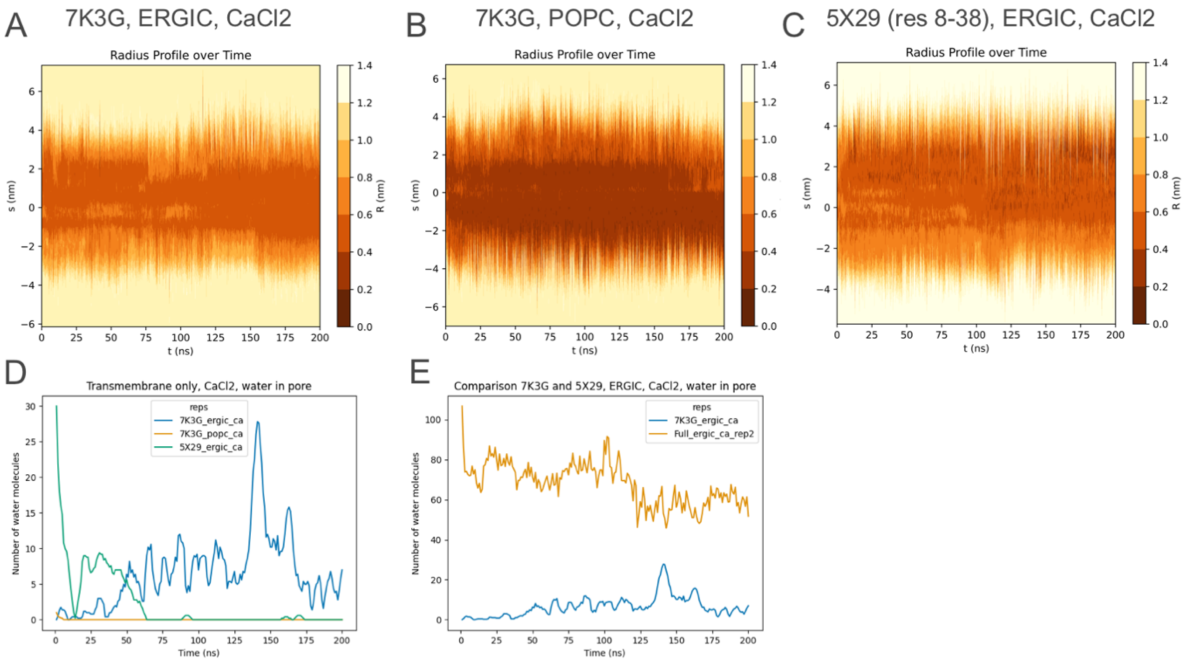

Figure S10: Radius profiles and hydration levels of ETM constructs used in our MD simulations in membranes.

Table S1: Interaction of POPI and POPS with the first four residues on the luminal face of SARS-CoV-2 E in ERGIC mimetic membranes (distance cut-off 3.5 Å, distance between any POPI or POPS and residues Glu8, Thr9, Gly10 and Glu11)

| Sars-CoV-2 E in ERGIC mimetic membrane with $CaCl_2$ | | | |
|---|---|---|---|
| **Rep** | **Interaction POPI – LUM (%)** | **Interaction POPS – LUM (%)** | **No interaction (%)** |
| 1 | 34.8 | 13.3 | 63.8 |
| 2 | 23.0 | 0.0 | 77.0 |
| 3 | 36.9 | 0.4 | 62.7 |
| 4 | 44.6 | 0.4 | 55.1 |
| 5 | 21.4 | 0.0 | 78.6 |
| Sars-CoV-2 E in ERGIC mimetic membrane with NaCl | | | |
| 1 | 33.4 | 0.0 | 66.6 |
| 2 | 36.7 | 0.0 | 63.3 |
| 3 | 44.1 | 0.0 | 55.8 |
| 4 | 23.8 | 0.0 | 76.2 |
| 5 | 14.8 | 0.0 | 85.2 |

Table S2: Interaction between palmitoyl chains and the hydrophobic gate Phe residues in the E protein in simulations of the E protein in ERGIC mimetic membrane with $Ca^{2+}$ ions (distance cut-off 3.5 Å, distance between atoms of Phe23 and Cys43)

| Sars-CoV-2 E in ERGIC mimetic membrane, with $CaCl_2$ | | | | | |
|---|---|---|---|---|---|
| **Rep** | **Chain** | **PHE – CYSP interaction** | **No interaction** | **PHE – CYSP interaction (%)** | **Average (%)** |
| 1 | 1 | 537 | 76 | 53.7 | 52.8 |
| | 2 | 607 | 6 | 60.7 | |
| | 3 | 554 | 59 | 55.4 | |
| | 4 | 596 | 17 | 59.6 | |
| | 5 | 347 | 266 | 34.7 | |
| 2 | 1 | 591 | 409 | 59.1 | 82.24 |
| | 2 | 981 | 19 | 98.1 | |
| | 3 | 923 | 77 | 92.3 | |
| | 4 | 976 | 24 | 97.6 | |
| | 5 | 641 | 359 | 64.1 | |
| 3 | 1 | 846 | 154 | 84.6 | 84.82 |
| | 2 | 554 | 446 | 55.4 | |
| | 3 | 993 | 7 | 99.3 | |
| | 4 | 962 | 38 | 96.2 | |
| | 5 | 886 | 114 | 88.6 | |
| 4 | 1 | 910 | 90 | 91 | 57.5 |
| | 2 | 883 | 117 | 88.3 | |
| | 3 | 484 | 516 | 48.4 | |
| | 4 | 383 | 617 | 38.3 | |
| | 5 | 215 | 785 | 21.5 | |
| 5 | 1 | 936 | 32 | 96.7 | 72.2 |
| | 2 | 539 | 429 | 55.7 | |
| | 3 | 414 | 554 | 42.8 | |
| | 4 | 657 | 311 | 67.9 | |
| | 5 | 949 | 19 | 98.0 | |

Table S3: Interaction between palmitoyl acyl chains and hydrophobic gate Phe residues in the E protein in simulations of the E protein in ERGIC mimetic membrane with $Na^+$ ions (distance cutoff 3.5 Å, distance between atoms of Phe23 and Cys43)

| Sars-CoV-2 E in ERGIC mimetic membrane, with NaCl | | | | | |
|---|---|---|---|---|---|
| **Rep** | **Chain** | **PHE – CYSP interaction** | **No interaction** | **PHE – CYSP interaction (%)** | **Average (%)** |
| 1 | 1 | 96 | 904 | 9.6 | 52.8 |
| | 2 | 1000 | 0 | 100 | |
| | 3 | 990 | 10 | 99 | |
| | 4 | 668 | 332 | 66.8 | |
| | 5 | 512 | 488 | 51.2 | |
| 2 | 1 | 291 | 709 | 29.1 | 81.3 |
| | 2 | 991 | 9 | 99.1 | |
| | 3 | 974 | 26 | 97.4 | |
| | 4 | 964 | 36 | 96.4 | |
| | 5 | 843 | 157 | 84.3 | |
| 3 | 1 | 1001 | 0 | 100.0 | 78.7 |
| | 2 | 823 | 178 | 82.2 | |
| | 3 | 601 | 400 | 60.0 | |
| | 4 | 712 | 289 | 71.1 | |
| | 5 | 802 | 199 | 80.1 | |
| 4 | 1 | 506 | 494 | 50.6 | 76.6 |
| | 2 | 930 | 70 | 93 | |
| | 3 | 667 | 333 | 66.7 | |
| | 4 | 823 | 177 | 82.3 | |
| | 5 | 904 | 96 | 90.4 | |
| 5 | 1 | 63 | 937 | 6.3 | 43.7 |
| | 2 | 987 | 13 | 98.7 | |
| | 3 | 355 | 645 | 35.5 | |
| | 4 | 717 | 283 | 71.7 | |
| | 5 | 64 | 936 | 6.4 | |

Table S4: Interaction between palmitoyl acyl chains and hydrophobic gate Phe residues in the E protein in simulations of the E protein in POPC membrane with $Ca^{2+}$ ions (distance cutoff 3.5 Å, distance between atoms of PHE- and CYSP)

| Sars-CoV-2 E in POPC membrane, with $CaCl_2$ | | | | | |
|---|---|---|---|---|---|
| **Rep** | **Chain** | **PHE – CYSP interaction** | **No interaction** | **PHE – CYSP interaction (%)** | **Average (%)** |
| 1 | 1 | 521 | 479 | 52.1 | 62.4 |
| | 2 | 982 | 18 | 98.2 | |
| | 3 | 41 | 959 | 4.1 | |
| | 4 | 653 | 347 | 65.3 | |
| | 5 | 924 | 76 | 92.4 | |
| 2 | 1 | 716 | 284 | 71.6 | 66.7 |
| | 2 | 252 | 748 | 25.2 | |
| | 3 | 955 | 45 | 95.5 | |
| | 4 | 891 | 109 | 89.1 | |
| | 5 | 522 | 478 | 52.2 | |
| 3 | 1 | 976 | 24 | 97.6 | 52.6 |
| | 2 | 982 | 18 | 98.2 | |
| | 3 | 117 | 883 | 11.7 | |
| | 4 | 266 | 734 | 26.6 | |
| | 5 | 289 | 711 | 28.9 | |
| 4 | 1 | 871 | 52 | 94.4 | 33.3 |
| | 2 | 91 | 832 | 9.9 | |
| | 3 | 189 | 734 | 20.5 | |
| | 4 | 13 | 910 | 1.4 | |
| | 5 | 371 | 552 | 40.2 | |
| 5 | 1 | 814 | 106 | 88.5 | 41.6 |
| | 2 | 403 | 517 | 43.8 | |
| | 3 | 121 | 799 | 13.2 | |
| | 4 | 241 | 679 | 26.2 | |
| | 5 | 335 | 585 | 36.4 | |

Table S5: Residence time of $Ca^{2+}$ and $Na^{+}$ in occupied Glu binding sites at the luminal E terminus

| **E-residue on protein chain:** | **ERGIC** | | **POPC** |
|---|---|---|---|
| | **$Ca^{2+}$** | **$Na^{+}$** | **$Ca^{2+}$** |
| **A** | 1.3 ± 0.23 ns | 0.3 ± 0.03 ns | 1.0 ± 0.11 ns |
| **B** | 1.3 ± 0.24 ns | 0.2 ± 0.01 ns | 1.4 ± 0.22 ns |
| **C** | 1.0 ± 0.16 ns | 0.3 ± 0.02 ns | 0.9 ± 0.08 ns |
| **D** | 1.3 ± 0.15 ns | 0.3 ± 0.01 ns | 1.0 ± 0.11 ns |
| **E** | 1.5 ± 0.25 ns | 0.3 ± 0.05 ns | 2.1 ± 0.6 ns |

Table S6: Probability of number of Glu binding sites at the luminal E terminus occupied concurrently

| **Number of E-residues** | **ERGIC** | | **POPC** |
|---|---|---|---|
| | **$Ca^{2+}$** | **$Na^{+}$** | **$Ca^{2+}$** |
| **0** | 0.39 ± 0.03 | 0.85 ± 0.01 | 0.47 ± 0.03 |
| **1** | 0.42 ± 0.01 | 0.14 ± 0.01 | 0.4 ± 0.02 |
| **2** | 0.16 ± 0.01 | 0.01 ± 0.0 | 0.11 ± 0.02 |
| **3** | 0.03 ± 0.0 | 0.0 ± 0.0 | 0.001 ± 0.0 |
| **4** | 0.0 ± 0.0 | 0.0 ± 0.0 | 0.0 ± 0.0 |
| **5** | 0.0 ± 0.0 | 0.0 ± 0.0 | 0.0 ± 0.0 |
